# Retinal electrical synapse plasticity is required for optimal visual performance

**DOI:** 10.64898/2026.07.27.741000

**Authors:** Ya-Ping Lin, Cheryl K. Mitchell, Nange Jin, Stephan Tetenborg, Nikki A. Brantley, Zhijing Zhang, Anna Naglis, Ye Long, Hongyan Li, Suzette Luke, Chai-An Mao, Steven W. Wang, Christophe P. Ribelayga, John O’Brien

## Abstract

Most electrical synapses in the mammalian central nervous system are composed of Connexin 36 (Cx36). Electrical synapses are functionally plastic, changing their degree of coupling based on the activity of the cell or connected cells, or based on activation of neurotransmitter or neurohormone receptors. Plasticity can reach an extreme in which electrical synapses become functionally silent, which is a normal operational condition for some circuits. Cx36 coupling is regulated by phosphorylation, which opens the channels. In retinal circuits, Cx36 is often maintained in a poorly phosphorylated, poorly coupled state. We reasoned that phosphomimetic mutants of Cx36 could remain constitutively open and maintain circuits in a well-coupled state that will interrogate the need for plasticity. We developed a constitutively open Cx36 mutant by systematically replacing phosphorylatable residues that regulate coupling with acidic residues. Single mutants of serine 315 significantly modified functional regulation of coupling in HeLa cells, but mutation of four residues was required to produce a mutant that was constitutively open. This mutant, Cx36-S110D, T111E, S293D, S315D, called Cx36-DEDD, displayed high coupling in control conditions and only modest changes under phosphorylating and dephosphorylating conditions. We developed a conditional knockin mouse that expresses Cx36-DEDD and cytoplasmic tdTomato in cells that expressed Cre recombinase. When crossed with Six3-Cre mice, Cx36-DEDD expressed widely in the retina including in photoreceptors, bipolar, amacrine and ganglion cells. Rod-cone electrical coupling displayed the maximum of its physiological dynamic range, and photopic visual acuity and contrast sensitivity were significantly reduced in Cx36-DEDD homozygous animals. We conclude that reduction of coupling in some retinal circuits is required for optimal daylight vision.

**Significance Statement:** Two types of synapses, chemical and electrical, work together throughout the central nervous system to perform neurological functions. While it is widely understood for chemical synapses that plasticity, changing the strength of synaptic connections, plays critical roles in many processes, this is far less understood for electrical synapses. By developing an electrical synapse protein mutant that locks channels in an open state, we have investigated retinal circuits that retain functional electrical synapses but lack their latitude for plasticity. This perturbation significantly compromises visual acuity and contrast sensitivity in the daylight, revealing that electrical synapse plasticity is necessary to tune retinal functions for optimal performance. Thus, electrical synapse plasticity along with chemical synapse plasticity is required for neural function.

## Introduction

Across almost all nervous systems studied to date, synapses are comprised of both chemical and electrical types. Electrical synapses, formed by gap junctions among neurons, serve functions complementary to those of chemical synapses. Chemical synapses generally form amplifying feedforward or feedback synapses within chains of neurons of different types, conveying signals of either the same or opposite sign that define circuit function. In contrast, electrical synapses are limited to transmitting sign-conserving signals at gains less than 1. In spite of these “limitations,” electrical synapses have profound effects on the circuits they subserve. When organized within groups of the same neuron type, electrical synapses share signals that can coordinate groups of neurons to function in synchrony. When organized between neurons of different type, electrical synapses can coordinate function of heterologous groups of neurons and form defined feedforward circuits with intrinsic capability for bidirectional signal propagation.

A fundamental property of nervous systems is dynamic plasticity of their synapses. Synaptic plasticity is central to learning and memory, adaptation to environmental cues, attention and many other aspects of nervous system function (1, 2). It has long been known that plasticity is not merely the domain of chemical synapses, but that electrical synapses also display functional plasticity (3–5). Electrical synapse plasticity modifies the size of groups of neurons that behave in a coordinated fashion (6). This can have effects ranging from subtle changes in spike synchrony that modifies the information coding transmitted to downstream neurons (7, 8) to complete reconfiguration of circuits (9–12). These changes are central elements of a nervous system’s ability to tune performance to meet the demands of an organism’s environmental experience.

In many electrically-coupled networks, it is clear that functional plasticity is achieved through changes in the state of the electrical synapse proteins without significant loss or increase in their abundance. In retinal horizontal cells, light adaptation and dopamine receptor signaling induce profound changes in functional coupling (3, 13–17). Similar interventions result in changes in the packing density and organization of horizontal cell gap junctions, but do not change the number of channels (18, 19), suggesting that some change in the functional state of the gap junction protein is responsible for the change in coupling. Similar to horizontal cells, mammalian retinal type 2 (AII) amacrine cells display large changes in functional coupling driven by dopamine receptor signaling (20, 21) and light adaptation (22, 23). Studies of Cx36 phosphorylation in AII amacrine cells treated with dopaminergic agonists and antagonists showed a direct, positive correlation of functional coupling to Connexin 36 (Cx36) phosphorylation state over a 20-fold dynamic range of coupling, with no changes in the number or size of gap junctions (24). This strongly suggests that Cx36 phosphorylation is the primary determinant of electrical synapse functional plasticity in the responses of the AII amacrine cell network to dopamine signaling. Photoreceptor networks are also subject to dramatic changes in gap junction coupling controlled by light, dopamine receptor signaling and the circadian clock (25–27). Similar to the situation in AII amacrine cells, a 20-fold change in tracer coupling was quantitatively and positively correlated to changes in Cx36 phosphorylation state, with no changes in the number of gap junctions (28, 29). Such studies implicate Cx36 phosphorylation as a primary point for regulation of functional plasticity of the electrical synapses that use this connexin.

Over the course of many studies examining the phosphorylation state of Cx36 gap junctions in various retinal networks in several species of animals, we have observed that the baseline state of most gap junctions is one with low phosphorylation. This implies that most electrical synapses support relatively low coupling, but have significant latitude for enhancement. Because of the significant roles that electrical synapse plasticity has in homeostatic regulation and adaptation of neural networks, we set out to develop an animal model in which electrical synapses are locked at the upper limit of their dynamic range for functional plasticity. This would allow investigators to understand the functional implications of electrical coupling at the upper limit of normal physiological regulation, and in so doing understand the contribution of electrical synapse plasticity to neural network function. In this study, we have identified the regulatory phosphorylation sites of mouse Cx36 that are required to open the channel and developed phosphomimetic mutants that imitate the phosphorylated state. We identified one mutant with four phosphorylation sites converted to phosphomimetic aspartate or glutamate residues that was locked in an open state and could not be uncoupled by protein kinase A signaling. We used this mutant to develop a conditional knockin mouse that expresses the mutant when crossed with a strain expressing Cre recombinase in a cell or tissue type of interest. When implemented in the retina, these changes reduced peak photopic visual acuity and contrast sensitivity, indicating that these functions of the retina require reducing electrical coupling in some circuits. This mouse line will be a useful tool to investigate the requirement for electrical synapse plasticity in neural networks.

## Results

### Phosphomimetic mutants of Cx36 constitutively open the channels

*In vitro* studies of Cx36 and homologues have extensively mapped phosphorylation sites and inferred their modes of action. In early studies, phosphorylation was thought to reduce coupling, as activating protein kinase A (PKA) typically led to less coupling in retinal neurons and expression systems. Mitropoulou and Bruzzone (30) identified a protein kinase A consensus recognition motif in perch Cx35, absent in skate Cx35, that targeted serine 110 (Ser110) in the cytoplasmic loop for phosphorylation (Fig. 1A and B). Disruption of this motif blocked the suppression of hemichannel currents caused by activation of PKA. Ouyang *et al*. (31) showed directly that this site, along with Ser276 (Ser293 in mammalian Cx36) in the C-terminal tail, was phosphorylated by PKA, and that mutation of these two sites to non-phosphorylatable alanine residues suppressed both negative and positive regulation of coupling by PKA signaling. The same two sites were also found to be phosphorylated by the nitric oxide-regulated protein kinase G (PKG) (32)(Fig. 1A and B), leading to comparable regulation of the channels. While both PKA and PKG signaling pathways led to reduced coupling in expression systems and some neurons, Kothmann *et al*. (24) showed that phosphorylation at these sites was strongly correlated with increased coupling. The disparity in observations was explained by the discovery that the dominant action of the PKA signaling pathway in the neuron studied was to activate protein phosphatase 2A (PP2A), which dephosphorylated Cx36 (24). This PP2A signaling pathway was later shown to be active in the Hela cell expression system used to study these connexins, revealing a signaling module shared between certain retinal neurons and the Hela cell expression system (33).

**Figure 1.**
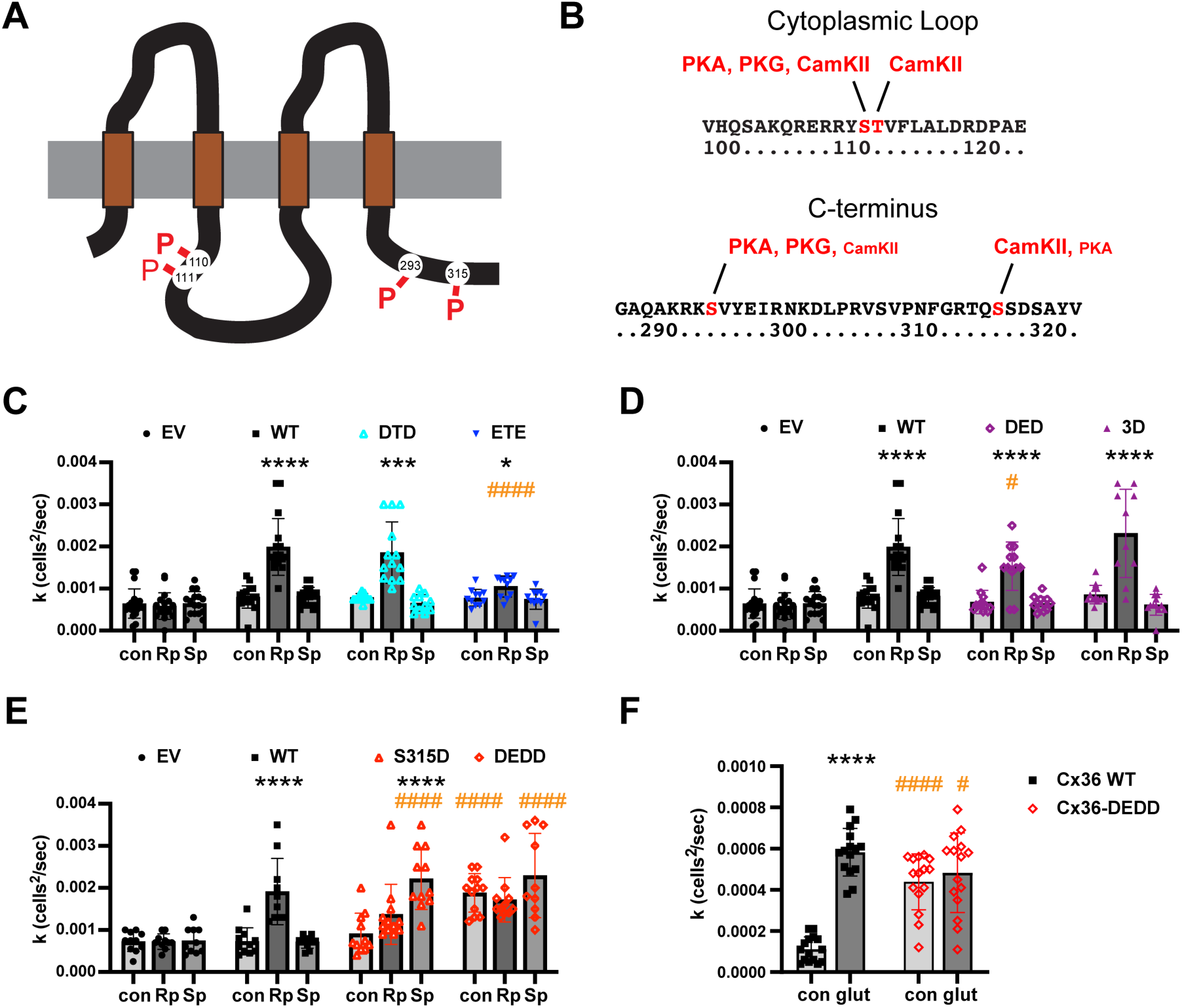
Connexin 36 phosphorylation sites and phosphomimetic mutants. **A.** Schematic diagram of Cx36 showing locations of phosphorylation sites in the middle cytoplasmic loop domain and carboxyl terminal domain. **B.** Amino acid sequence of mouse Cx36 in the region of the phosphorylation sites in the middle cytoplasmic loop (top) and C-terminal domain (bottom). Phosphorylated residues are illustrated in red font, and protein kinases known to phosphorylate those sites are displayed. Kinases that are observed to phosphorylate a site weakly in *in vitro* studies are shown in reduced size font. **C.** Tracer coupling studies of mouse Cx36-EGFP constructs transfected in Hela cells. Diffusion coefficients (k) for Neurobiotin transfer are shown in control medium without drugs (con), with 10 µM PKA inhibitor (Rp) and 10 µM PKA activator (Sp). Constructs shown are wild type Cx36 (WT), Cx36-S110D, S293D (DTD), Cx36-S110E, S293E (ETE) and empty vector control (EV) expressing cytoplasmic EGFP. **D.** Tracer coupling studies of mouse Cx36 constructs WT, Cx36-S110D, T111E, S293D (DED), Cx36-S110D, T111D, S293D (3D), and empty vector control (EV). **E.** Tracer coupling studies of mouse Cx36 constructs WT, Cx36-S315D (315D), Cx36-S110D, T111E, S293D, S315D (DEDD), and empty vector control (EV). **F**. Tracer coupling studies of mouse Cx36 WT and Cx36-DEDD in no-drug control medium (con) or with 100 µM glutamate + 100 µM glycine (glut). Bars are means +/− SD; Bars include 10-20 measurements from 2-3 separate experiments. EV and WT controls in C-E were performed in the same experimental sessions as the mutants tested. Some EV and WT data are replicated in C-E. Two-way ANOVA with Tukey’s multiple comparison tests (C-E) or Fisher’s Least Significant Difference (F). * represent comparisons to the no drug control condition within each construct; # represent comparisons to the equivalent drug condition of Cx36 WT. * or #, p<0.05; ** or ##, p<0.01; *** or ###, p<0.001; **** or ####, p<0.0001.

Ouyang *et al*. (31) also found PKA to phosphorylate Ser298 (Ser315 of mammalian Cx36) weakly (Fig. 1A and B). This site is more potently phosphorylated by calmodulin-dependent protein kinase II (CamKII) (34), which leads to increased coupling in Cx36 (35). Alev *et al*. (34) found CamKII to phosphorylate both Ser110 and the adjacent Thr111 in the cytoplasmic loop domain (Fig. 1A and B). The latter residue is not phosphorylated by either PKA or PKG. These studies together identified four phosphorylation sites that regulated Cx36 coupling, with convergence of several protein kinase signaling pathways on these same residues (Fig. 1A and B).

Because studies that directly compared Cx36 phosphorylation with coupling found that phosphorylation increases coupling (24, 28), we reasoned that introducing mutations that mimic the phosphorylated state could increase coupling of Cx36 gap junctions or prevent them from being closed completely. To investigate this hypothesis, we systematically introduced phosphomimetic mutations into mouse Cx36 (36) and tested functional regulation of coupling in transfected Hela cells. We mutated the phosphorylatable serine or threonine residues to aspartic acid (Asp or D) or glutamic acid (Glu or E), each of which carry a negative charge on the side chain at neutral pH. In the first round of experiments, we mutated Ser110 and Ser293, the two residues identified to account for the predominant functional regulation (31), to Asp or Glu. We examined tracer coupling in transfected Hela cells in a paradigm in which cells were treated with the membrane-permeant PKA inhibitor Rp-8-cpt-cAMPS – 10 µM, or the membrane-permeant PKA activator Sp-8-cpt-cAMPS – 10 µM. In this paradigm, inhibition of PKA significantly increased tracer coupling of wild type (WT) Cx36 (Two-way ANOVA with Tukey’s multiple comparisons; p < 0.0001; statistics details in supplemental Table 1) while PKA activation had no measurable effect (Fig. 1C; suppl. Table 1). As alluded to above, activation of PKA in this system activates the phosphatase PP2A to reduce Cx36 phosphorylation and coupling, while inhibition of PKA suppresses PP2A activity, allowing Cx36 to be phosphorylated by an as yet unidentified protein kinase.

Hela cells have some background connexin expression, which is revealed in tracer coupling experiments with empty vector (EV) control cells transfected with EGFP vector. In EV controls, there was no measurable effect of PKA inhibitor (Rp) or PKA activator (Sp) treatments (Fig. 1C; suppl. Table 1). Furthermore, tracer coupling in WT Cx36 was statistically indistinguishable from EV controls in both control (no drug) and Sp conditions (Two-way ANOVA with Tukey’s multiple comparisons; Suppl. Table 1), indicating that Cx36 channels are largely closed in this assay system in those two conditions. Cx36 clones incorporating the mutations Ser110Asp and Ser293Asp (abbreviated DTD) displayed essentially normal regulation of coupling, with Rp significantly increasing coupling vs. control and no conditions significantly different from the equivalent conditions in WT Cx36 (Fig. 1C; suppl. Table 1). Cx36 clones incorporating the mutations Ser110Glu and Ser293Glu (abbreviated ETE) displayed suppressed, but still statistically significant enhancement of coupling during PKA inhibition (Rp) and significantly less coupling than WT Cx36 in this condition (Fig. 1C; suppl. Table 1). This suggests that the Glu mutations compromised the ability of Cx36 to open in PKA inhibited conditions and that the Asp mutations at two sites, Ser110 and Ser293, were insufficient to significantly open the Cx36 channels.

It was surprising that mutations of Ser110 and Ser293 did not cause significant opening of the channels, since mutation of those sites to non-phosphorylatable alanine residues had essentially eliminated regulation of coupling in the perch homologue of Cx36 (31). Nonetheless, there are other sites that are phosphorylated in Cx36, and these may have significant contributions to its opening. CamKII has been specifically identified to potently open gap junctions containing Cx36 and its homologues (35, 37, 38). Because CamKII uniquely phosphorylates Thr111 (Fig. 1B) in addition to the adjacent Ser110 (34), we next mutated this residue to either Glu or Asp in the Cx36-S110D, S293D (DTD) mutant. Both mutants, Cx36-S110D, T111E, S293D (abbreviated DED) and Cx36-S110D, T111D, S293D (abbreviated 3D) displayed significantly increased coupling when treated with the PKA inhibitor and displayed tracer coupling that was not significantly different from WT Cx36 in any of the drug treatment conditions (Fig. 1D; suppl. Table 1). Thus, a combination of three phosphomimetic mutations was still insufficient to produce constitutively open Cx36.

Following this round of mutations, a single residue known to regulate Cx36 coupling remained (Fig. 1A and B). Phosphorylation of Ser315 near the tip of the C-terminus by CamKII has been found to be responsible for the “run-up” in junctional coupling during whole-cell patch clamp experiments with cells expressing Cx36 (35), and phosphorylation at this residue alters which proteins are able to bind to the C-terminus of Cx36 (39, 40). Mutation of Ser315 by itself to phosphomimetic Asp caused a significant alteration in the pattern of regulation during inhibition or activation of PKA (Fig. 1E; suppl. Table 1). Both inhibition (Rp) and activation (Sp) of PKA significantly increased Cx36 coupling relative to control conditions, and coupling of Cx36-S315D was significantly higher than that of WT Cx36 during PKA activation. The functional “switching” of the effect of PKA activation on Cx36 coupling in the S315D mutant is, ironically, similar to the functional switching of a truncation mutant of Cx35 that specifically eliminates that phosphorylation site (31). This may be a reflection of the complex interplay between Cx36 and the proteins that associate with it and regulate its function (41).

As a last modification, we incorporated the S315D mutant into a Cx36 mutant with all of the other phosphorylation sites phosphomimetic: Cx36-S110D, T111E, S293D. We tested the resulting saturated mutant, abbreviated Cx36-DEDD, in the standard set of conditions manipulating PKA activity. Unlike all other mutants of Cx36 that we tested, Cx36-DEDD displayed significantly increased coupling in the control condition compared to WT Cx36 (Fig. 1E; suppl. Table 1), resisting the PKA-dependent dephosphorylating drive of PP2A activity that is constitutively active in this expression system. Coupling in the PKA activating condition (Sp) was also significantly higher than in WT Cx36. Finally, neither PKA inhibiting (Rp) nor PKA activating (Sp) condition was significantly different than the control condition for the Cx36-DEDD mutant (Fig. 1E; suppl. Table 1), indicating that these channels were constitutively open and were minimally regulated by PKA activity. To confirm the constitutively open nature of the Cx36-DEDD mutant, we tested it in Hela cells treated with 100 µM glutamate + 100 µM glycine, which activates endogenous NMDA receptors and subsequently CaMKII (42). While glutamate stimulation significantly enhanced coupling in WT Cx36, it did not change coupling in Cx36-DEDD and Cx36-DEDD coupling in control condition was significantly higher than WT Cx36 (Fig. 1F; suppl. Table 1). Thus, the Cx36-DEDD mutant remains persistently in the maximally coupled state achievable in the Hela cell expression system.

In order to make Cx36 channels constitutively open, it was necessary to make *all* known phosphorylation sites phosphomimetic. This provides insight into the functional regulation of Cx36 channels, implying that only fully phosphorylated channels open to contribute to electrical synapse function. This is perhaps not surprising given that electrical synapses comprised of Cx36 and its homologues become functionally silent at times, but it establishes a demanding condition that in order to open all channels in a gap junction, essentially all Cx36 subunits must be fully phosphorylated.

### Development of a Cx36 phosphomimetic mutant mouse

Cx36 knockout mice have been invaluable in studies of neuronal circuit functions.

The earliest studies of two independent Cx36 knockout mouse lines identified a major role for Cx36 in supporting oscillatory behavior of cortical and hippocampal networks (43–45) and an essential role in signal transmission in the retinal rod pathways (46, 47). Many other studies using these animals have revealed a large number of roles for Cx36 in circuits throughout the central nervous system. However, the large dynamic range for plasticity of circuits coupled by Cx36 (20, 21, 23, 24, 27) and the observation that most Cx36 gap junctions in some tissues exist in a poorly phosphorylated and hence poorly coupled state (24) suggests that weak coupling is the norm for many circuits. Thus, a Cx36 knockout mouse does not provide a full understanding of the influence of electrical synapses on neural circuit function. To provide an animal model that will allow investigators to understand the influence of electrical coupling at its functional limits and to understand the roles of electrical synapse plasticity in neural circuits, we set out to develop a Cx36 phosphomimetic knockin mouse.

We used the saturated phosphomimetic mutant, Cx36-S110D, T111E, S293D, S315D (Cx36-DEDD) to develop targeting constructs for gene knock-in via homologous recombination. All of the functional mutations are contained within the second exon of the *Cx36* (*gjd2*) gene (e.g. Fig. 2A) so this exon was targeted for replacement with a mutant-containing construct. Our initial strategy was to replace wild type exon 2 with the mutant exon and a marker gene. While we were able to isolate correctly targeted embryonic stem cell (ES cell) lines, we were unable to obtain knock-in mice. In 3 experiments with 2 different correctly targeted ES cell clones, chimeric F0 pups were born but did not survive. The pups either died postnatally or were eaten by their parents, implying that the Cx36-DEDD protein disrupted normal brain function.

**Figure 2.**
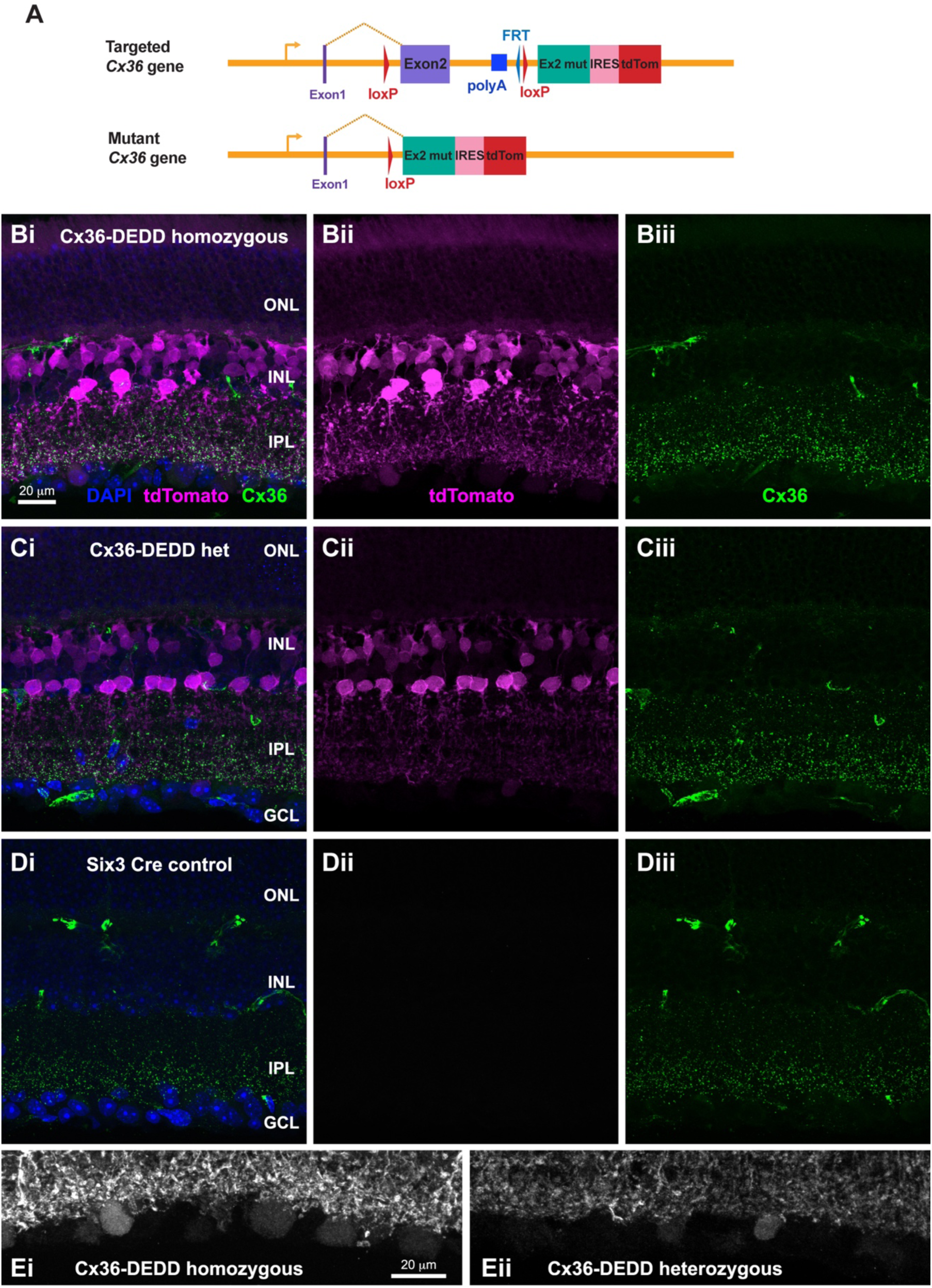
Cx36-DEDD knock-in mouse. **A.** Connexin 36 (*gjd2*) gene targeted for conditional knock-in of phosphomimetic Cx36-DEDD second exon. Wild type protein-coding exons 1 and 2 are shown with purple color; mRNA transcript initiates at the gold right arrow and exons are spliced as shown by the gold chevron. The “wild type” transcript terminates at an artificial polyadenylation signal (“polyA” – blue) after exon 2. Exon 2 and the artificial polyA signal are flanked by loxP signals (red) and followed by a cassette containing the DEDD mutant exon 2, an internal ribosome entry sequence (IRES) and tdTomato coding sequence. Frt site is a remnant of a self-excising antibiotic resistance gene cassette used to select for targeted embryonic stem cell lines. When wild type exon 2 is deleted by expression of Cre recombinase in cells of interest (second line), the transcript splices into the mutant exon 2 and terminates at natural polyadenylation sequences in the Cx36 gene. **Bi.** Immunostaining of retina from Cx36^(DEDD/DEDD)^/Six3-Cre mouse. Nuclear layers stained with DAPI (blue) are the outer nuclear layer (ONL), inner nuclear layer (INL) and ganglion cell layer (GCL; only shown in Ci and Di); inner plexiform layer (IPL) is also labeled. Cx36 is labeled with anti-Cx36 antibody (green) and tdTomato signal is boosted with anti-RFP antibody (magenta). **Bii**. tdTomato signal alone. **Biii**. Cx36 signal alone. Irregular signals at outer and inner edges of the INL are non-specific labeling of capillaries by the anti-mouse secondary antibody. **Ci-Ciii**. Immunostaining of retina from Cx36^(DEDD/+)^/Six3-Cre mouse. Color channels, labels and panel order are as in B. **Di-Diii**. Immunostaining of retina from Six3-Cre transgenic mouse in wild type background. Color channels, labels and panel order are as in B. Scale bar in Bi applies to all B-D panels. **Ei**. tdTomato signal in ganglion cell layer from Cx36^(DEDD/DEDD)^/Six3-Cre mouse (panel Bii) with brightness doubled. **Eii**. tdTomato signal in ganglion cell layer from Cx36^(DEDD/+)^/Six3-Cre mouse (panel Cii) with brightness doubled. Scale bar in Ei applies to Eii as well.

Because of our failure to obtain mice with a direct knock-in approach, we chose to develop a conditional knock-in construct that would express Cx36-DEDD only in cells in which Cre recombinase had been expressed. Figure 2A shows a diagram of the Cx36 gene with mutant elements knocked in. In this construct, the wild type exon 2 is followed by an artificial polyadenylation signal and flanked by loxP sites. These elements are followed by the DEDD mutant exon 2, an internal ribosome entry sequence (IRES) and the coding sequence of tdTomato. Expression of the complete knock-in construct (Fig. 2A top line – mRNA transcript initiated at the right arrow and spliced as indicated by the gold chevron) produces an mRNA that contains the wild type exon 1 and 2 and terminates at an artificial polyadenylation signal. This transcript will express the wild type Cx36 protein, but will lose some elements that may regulate the mRNA transcript through interactions with the very long 3’ untranslated region (36), since only the proximal 1 kb of 3’ untranslated region is retained.

When crossed with a suitable Cre recombinase expressing mouse line or when Cre is delivered via viral transduction, the wild type Cx36 exon 2 and artificial polyA signal are deleted (Fig. 2A second line). The mRNA transcript in this case splices into the DEDD mutant exon 2 and continues through the IRES and tdTomato sequences and into the natural 3’ untranslated region of the Cx36 gene. This single transcript expresses the Cx36-DEDD mutant protein with a natural C-terminus (not blocked by a fluorescent protein or residues of self-cleaving sequences) and, separately, soluble tdTomato.

Figure 2B shows a section of retina from a Cx36^(DEDD/DEDD)^ mouse expressing a Six3-Cre transgene (48), which expresses in retinal progenitor cells, and so releases Cx36-DEDD broadly throughout the retina. Cx36 labeling was present in an expected distribution in the retinal inner plexiform layer (IPL) and outer plexiform layer (Figure 2Bi, Biii). We amplified tdTomato signal with an anti-red fluorescent protein antibody. Labeling could be detected readily in amacrine and bipolar cell somata in the inner nuclear layer (INL), faintly in photoreceptors, and very faintly in retinal ganglion cells (Figure 2Bi, Bii). Figure 2C shows the same labeling in Cx36^(DEDD/+)^ heterozygous mouse expressing the Six3-Cre transgene, revealing similar labeling patterns for both tdTomato and Cx36. Figure 2D shows the same labeling scheme in a control mouse expressing the Six3-Cre transgene but not containing the Cx36-DEDD knock-in allele. Cx36 expression in this retina follows the wild type pattern and is essentially the same as in the Cx36-DEDD knock-in mice. We noted that tdTomato signal in retinal ganglion cells was extremely faint at imaging settings that revealed cells in the inner nuclear layer well. Figure 2E shows the tdTomato signal in the retinal ganglion cell layer at twice the brightness shown in panels B-D. With this signal enhancement, one can see expression of tdTomato in a number of retinal ganglion cells in both the Six3-Cre/Cx36^(DEDD/DEDD)^ homozygous mouse (Figure 2Ei) and the Six3-Cre/Cx36^(DEDD/+)^ heterozygous mouse (Figure 2Eii). This is in agreement with the known expression of Cx36 in some types of retinal ganglion cells (49–53), including the intrinsically photosensitive retinal ganglion cell (ipRGC) (54), but suggests that the expression level of the gene is very low in these cell types. The marker gene expression in retinal neurons suggests that the Cx36-DEDD mutant gene is expressed in these mice.

In both homozygous and heterozygous Cx36-DEDD mice, a row of cells at the bottom of the INL showed the brightest tdTomato signal (e.g. Figure 2Cii). These cells have the characteristics of AII amacrine cells, with lobular appendages in the Off sublamina of the IPL and branching dendrites in the On sublamina of the IPL that extensively colocalize with Cx36 puncta. To verify that these are AII amacrine cells, we injected eyes of Six3-Cre/Cx36^(DEDD/DEDD)^ homozygous mice with an adeno-associated virus expressing EGFP driven by the synthetic HKamac promoter that has previously been shown to express in AII amacrine cells (55). Figure 3 shows that EGFP expression colocalizes with tdTomato expression in cells labeled by the virus. Furthermore, these cells have large numbers of Cx36 gap junctions on their arboreal dendrites where they make contact with other amacrine cells as well as along the dendrite shafts where they do not make contacts with other cells of the same type (Figure 3D). These are likely to be gap junctions onto cone bipolar cells, a prominent feature of AII amacrine cells.

**Figure 3.**
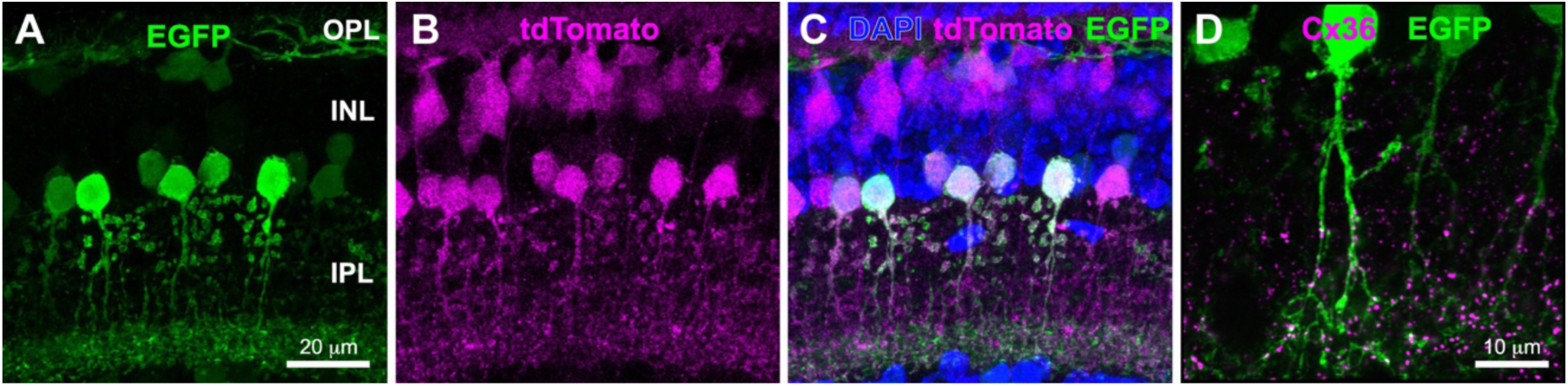
Cx36-DEDD expression in AII amacrine cells. **A.** Expression of EGFP driven by the HKamac promoter in cells transduced by an Adeno-associated virus. The HKamac promoter labels predominantly AII amacrine cells (INL and IPL) and horizontal cells (INL and OPL), with sparse labeling of another amacrine cell type. **B.** Expression of tdTomato in a Cx36^(DEDD/DEDD)^/Six3-Cre mouse. **C.** Merged image shows that the major cell type with strong tdTomato expression colocalizes with EGFP driven by the HKamac promoter, identifying them as AII amacrine cells. OPL – outer plexiform layer, INL – inner nuclear layer, IPL – inner plexiform layer. **D.** HKamac-EGFP labeled AII amacrine cells in WT mouse retina co-localize extensively with immunolabeling for Cx36.

Thus, these cells can be confidently identified as AII amacrine cells.

### Cx36-DEDD expression increased neuronal gap junction coupling

The goal of developing the Cx36-DEDD mouse was to be able to lock Cx36 gap junctions in an open state so that the maximum of the dynamic range of expected electrical synapse plasticity could be investigated. This unique limit state also allows the investigator to assess what is the role of electrical synapse plasticity in neural circuits of interest. To determine whether we have accomplished this, we recorded from rod-cone pairs in isolated retina from Six3-Cre/Cx36^(DEDD/DEDD)^ homozygous mice and from Six3-Cre control mice. Figure 4A shows voltage clamp recordings of rod-cone pairs from a control mouse and a Cx36-DEDD homozygous mouse. Junctional currents were substantially higher in the Cx36-DEDD homozygous rod-cone pair. The current voltage relationships of three Cx36-DEDD homozygous and two Six3-Cre control rod-cone pairs are shown in Figure 4B. The steeper slope of the Cx36-DEDD current-voltage relationship indicates higher junctional conductance. Individual junctional conductances of measured rod-cone pairs are plotted in the histogram in Figure 4C, along with the means (color-coded dashed lines). The mean conductance of homozygous Cx36-DEDD rod-cone pairs was 1302 ± 251 pS, significantly higher than the 375 ± 206 pS mean conductance of Six3-Cre rod-cone pairs (Welch’s T-test; p = 0.026). To put this conductance in context, we compared these values to those of C57Bl6 mice in the same control condition and with treatments to activate or inhibit D2-like dopamine receptors reported by Jin et al. (27). While C57Bl6 mice in control conditions exhibited a very similar mean conductance (453 pS) to our control group, activation of dopamine D2-like receptors with quinpirole, mimicking a light-adapted state, reduced rod-cone junctional conductance to 19 pS (Fig. 4C). In contrast, inhibition of D2-like receptors, mimicking a nighttime, dark-adapted state, increased junctional conductance to 1070 pS. This is similar to the 1302 pS value we measured in Cx36-DEDD homozygous rod-cone pairs, suggesting these are in a maximally-coupled state.

**Figure 4.**
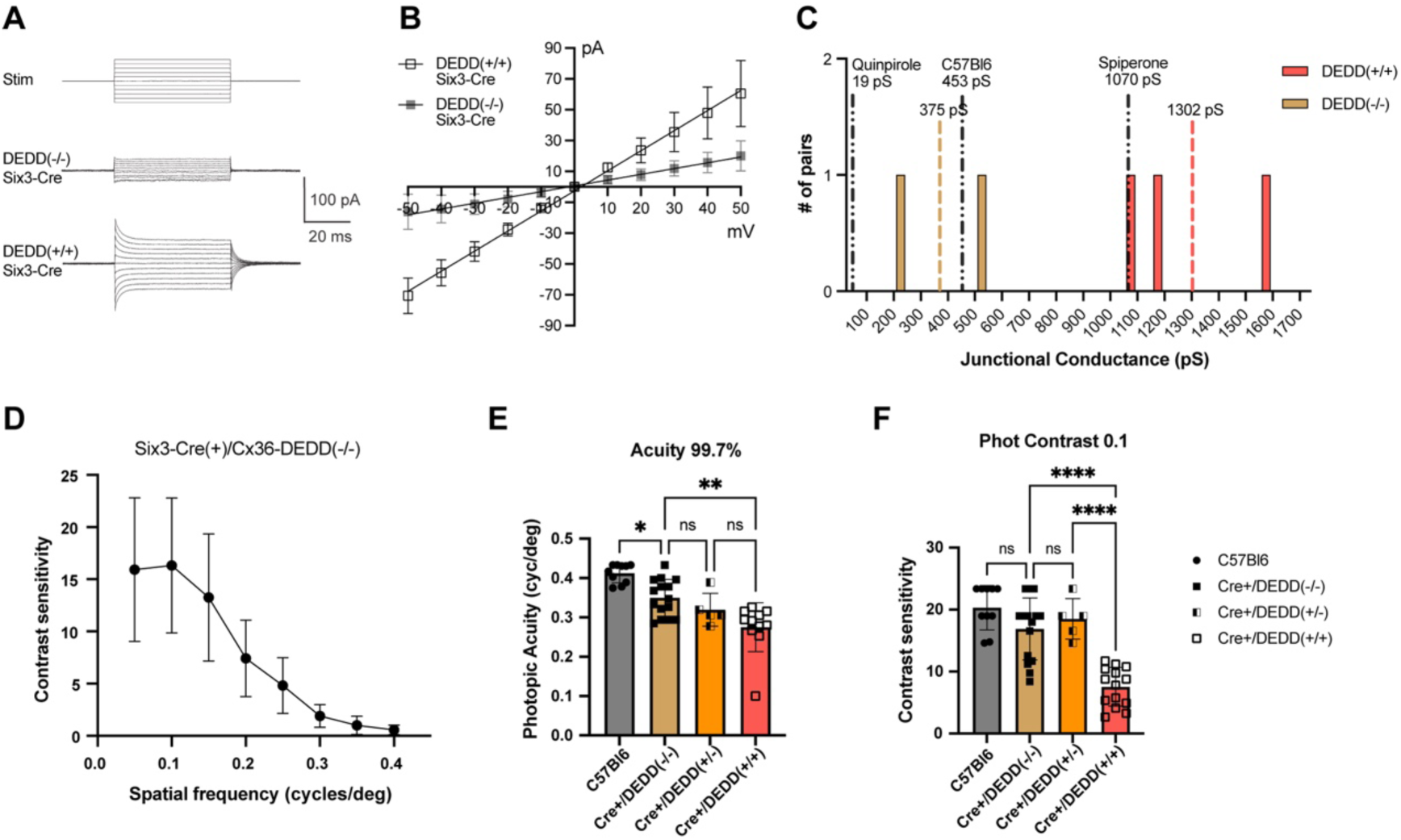
Retinal electrical coupling and photopic optomotor responses of Cx36-DEDD mice. **A.** Junctional currents between rod-cone pairs in response to voltage steps (top line) in 10 mV increments from −50 mV to +50 mV in a Six3-Cre control mouse and a Six3-Cre/Cx36^(DEDD/DEDD)^ homozygous mouse. **B**. Current-voltage relationships of rod-cone junctional currents from two Six3-Cre control mice and three Six3-Cre/Cx36^(DEDD/DEDD)^ homozygous mice. **C**. Histogram of rod-cone junctional conductance (measured from the slope of the current-voltage relationship) from pairs shown in B. The mean junctional conductance is shown with the color-coded dashed lines. Black dashed lines show mean conductances of rod-cone pairs from C57Bl6 mice in control condition and with treatments with quinpirole to close gap junctions and spiperone to open gap junctions from Jin et al. (27). **D.** Relationship between contrast sensitivity and visual acuity under photopic lighting conditions in Six3-Cre control mice. Data are means ± 1SD, n = 3 to 8 animals per data point. **E**. Pooled photopic visual acuity threshold measured at 99.7% contrast in C57Bl6 (control), Six3-Cre (control), Six3-Cre/Cx36^(DEDD/+)^ heterozygous and Six3-Cre/Cx36^(DEDD/DEDD)^ homozygous mice. **F.** Pooled photopic contrast sensitivity threshold in the four strains of mice shown in B. Bars are means +/− SD. One-way ANOVA with Tukey’s multiple comparisons. ns = not significant; *, p<0.05; **, p<0.01; ***, p<0.001; ****, p<0.0001.

As a further test of neuronal coupling, we performed Neurobiotin tracer coupling experiments in AII amacrine cells of Cx36-DEDD mice crossed with Six3-Cre transgenic mice. Iontophoretic injections of Neurobiotin into single AII amacrine cells in whole mount retina led to tracer diffusion through a network of coupled AII amacrine and bipolar cells. Supplemental figure 1A and B show that an injected patch imaged at the level of AII amacrine cells in retina of a Six3-Cre/Cx36^(DEDD/DEDD)^ homozygous mouse displayed much more extensive tracer diffusion than a comparable patch in a Six3-Cre/Cx36^(DEDD/+)^ heterozygous mouse. The steady decrease in tracer labeling intensity with distance from the injected cell allows us to measure the diffusion coefficient for tracer transfer using a compartmental diffusion model (56). Tracer injections into AII amacrine cells in Six3-Cre/Cx36^(DEDD/+)^ heterozygous mice yielded a mean diffusion coefficient *k* of 0.00013 ± 0.00004 cells^2^/sec (n = 6 injections in 3 animals). Injections into Six3-Cre/Cx36^(DEDD/DEDD)^ homozygous mouse AII amacrine cells yielded a mean diffusion coefficient *k* of 0.00049 ± 0.00015 cells^2^/sec (n = 5 injections in 3 animals), which was significantly higher (p = 0.0045; Welch’s t test). Thus, phosphomimetic Cx36 expressed homozygously significantly enhanced neuronal coupling over its heterozygous expression.

### Cx36-DEDD expression reduced photopic visual acuity and contrast sensitivity

Adaptation to photopic light triggers reductions in electrical coupling of several retinal neural networks. For example, while AII amacrine cells are capable of forming large coupled networks, the extent of coupling is reduced dramatically in light-adapted conditions (22, 23). This change in circuitry reduces pooling of signal passing through the rod pathway and enhances local crossover inhibition of the Off pathways by On pathway bipolar cells (57). Likewise, rod-cone coupling is reduced by light adaptation, dramatically suppressing rod input into the cone pathways (10, 25). Because Cx36-DEDD expression causes electrical synapses to remain locked in an open condition, we would predict that the loss of electrical synapse plasticity may compromise the functional shift from dark-adapted to light-adapted state. As a first test of this hypothesis, we examined the photopic optomotor response (OMR) to rotating vertical stripe patterns in Cx36-DEDD mice. The experiments systematically varied both stripe width and contrast (see methods) to establish visual acuity and contrast thresholds, respectively. Figure 4D shows the relationship between contrast sensitivity and visual acuity in photopic light in one of the control strains, Six3-Cre transgenic mice. Contrast sensitivity peaked at approximately 0.1 cycles/degree. This spatial frequency was used for subsequent tests of contrast sensitivity, while the maximum contrast produced in our system, 99.7 %, was used for tests of visual acuity.

We examined whether there were differences attributable to sex in the OMR assessments. Supplemental figure 2 shows that there were no differences between female and male mice in visual acuity tests in any genotype tested (Two-way ANOVA with Tukey’s multiple comparisons test; see Supplemental table 2 for full statistics details). Therefore, we combined female and male data in subsequent analyses.

Figures 4E and F show visual acuity and contrast sensitivity measures for Six3-Cre/Cx36^(DEDD/+)^ heterozygous mice and Six3-Cre/Cx36^(DEDD/DEDD)^ homozygous mice, as well as two control populations: wild type C57Bl6 mice and Six3-Cre transgenic mice (Cre+/DEDD-). Figure 4E shows that photopic visual acuity was slightly but significantly lower in Six3-Cre transgenic mice than in C57/Bl6 mice (One-way ANOVA with Tukey’s multiple comparisons test; Suppl. Table 2). Because of this difference, we used the Six3-Cre transgenic mice as the control population with which to compare Cx36-DEDD knock-in mice. Photopic visual acuity was significantly lower in Six3-Cre/Cx36^(DEDD/DEDD)^ homozygous mice than in Six3-Cre transgenic mice, while Six3-Cre/Cx36^(DEDD/−)^ heterozygous mice were not significantly different (Fig. 4E; Suppl. Table 2). In photopic contrast sensitivity tests, once again Six3-Cre/Cx36^(DEDD/DEDD)^ homozygous mice displayed significantly reduced contrast sensitivity than Six3-Cre transgenic control mice, while Six3-Cre/Cx36^(DEDD/−)^ heterozygous mice were not significantly different (Fig. 4F; Suppl. Table 2). Thus, expression of phosphomimetic Cx36 locked in an open configuration significantly compromised visual acuity and contrast sensitivity in photopic conditions.

## Discussion

Electrical synapse plasticity is a core element of central nervous system function, with changes in electrical coupling making key contributions to a wide variety of dynamic functional changes in neural networks (5, 58, 59). The value of Cx36 knockout mice to the study of electrical synapses in the mammalian central nervous system cannot be understated. However, loss of the protein only reveals one aspect of the multifaceted functionality of electrical synapses. In this study we have developed a mouse model that allows researchers to investigate another fundamental aspect of electrical synapse function: their dynamic plasticity. The Cx36-DEDD mouse encodes expression of a Cx36 mutant that is maximally open and unregulated, and can be investigated selectively in circuits of choice by cell type-specific expression of Cre recombinase. This allows us to investigate the limit state for specific electrical synapses and evaluate how circuits behave when this limit state is reached.

Our examination of the functional limits of coupling was done in the electrically coupled vertebrate photoreceptor network. Jin et al. estimated a 24-fold difference in rod-cone junctional conductance between its minimum state in the presence of D2/D4 dopamine receptor agonist quinpirole and its maximum state in the presence of D2/D4 receptor antagonist spiperone (27). Using an estimated Cx36 single channel conductance of 14.3 pS (60, 61), and their estimates from 3D EM volumes of the size and number of gap junctions between rods and cones in mouse retina, Ishibashi et al. (62) calculated a theoretical maximum rod-cone conductance of 1228 ± 120 pS. This was considered statistically indistinguishable from the maximum conductance measured by Jin et al. (27) in the presence of spiperone, implying that approximately 100% of Cx36 channels were open in this condition. Within experimental error, the 1302 ± 251 pS conductance of rod-cone pairs in Six3-Cre/Cx36^(DEDD/DEDD)^ mice is indistinguishable from this value, implying that gap junctions comprised of Cx36-DEDD channels behave as if all channels are open.

The 24-fold dynamic range of mouse rod-cone junctional conductance measured by Jin et al. (27) is remarkably similar to the approximately 20-fold change in diffusion coefficient for tracer coupling through photoreceptor networks between daytime, light-adapted state and nighttime, dark-adapted state measured in both zebrafish and mouse retinas (28, 29), revealing a striking correlation between a quantitative measure of small molecule diffusion through the gap junctions and the electrical conductance of the gap junctions. The caveat of these measurements is that the minimum state for both types of measurements is so close to zero as to be nearly indistinguishable from the noise level. This situation further emphasizes that functional plasticity of the electrical synapse due to Cx36 phosphorylation governs a very large range of synapse activity, more than an order of magnitude. At the lower limit, electrical synapses can be essentially silent.

This feature of functional regulation is critical for reconfiguration of the rod circuits in the mammalian retina during the daytime (59), and is also a well-known feature of sensory-motor integration in the Mauthner cell circuit (63).

The most revealing finding of this study is that homozygous expression of Cx36-DEDD in most of the retina profoundly compromised photopic contrast sensitivity. The expression of Cx36-DEDD in photoreceptors and AII amacrine cells is expected to maintain the retina in a state that resembles the nighttime, dark-adapted state in which the gap junction elements of the primary and secondary rod pathways are maximally open and transmitting rod pathway signals. With this arrangement, one would expect that rod photoreceptors saturated by photopic lighting conditions would contribute signaling that compromises cone function. The prominent loss of contrast sensitivity in Cx36-DEDD homozygotes likely reflects this mechanism. However, the retina does have mechanisms that mitigate the effects of rod saturation in photopic conditions. First, it has been shown recently that mammalian rods escape saturation with prolonged adaptation to photopic conditions, contributing responses to contrast throughout the photopic range (64). This would enable primary and secondary rod pathways to encode contrast presented by the grating stimuli and transmit them to retinal ganglion cells. Second, some temporal contrast derived from rod photoreceptors is carried through a pathway independent of Cx36 (the tertiary rod pathway) (65). Thus, rod pathways do normally contribute to retinal ganglion cell responses in photopic conditions. Nonetheless, contrast coding was severely compromised by excessive coupling.

Photopic visual acuity was also compromised in homozygous Cx36-DEDD mice. Visual acuity depends on spatial tuning of the inhibitory surrounds of retinal ganglion cells. In primates, some ganglion cell spatial tuning may be inherited from bipolar cell receptive field surrounds imposed in the outer retina by horizontal cells (66). Horizontal cells are also coupled by electrical synapses, and their profound plasticity during light adaptation contributes significantly to temporal tuning of photoreceptor responses and the optomotor behavioral response (12). However, loss of gap junctions from horizontal cells, which use Cx57, did not alter retinal ganglion cell spatial response properties (67), suggesting an inner retinal origin for spatial response regulation. In contrast, loss of Cx36 throughout the retina reduced ganglion cell antagonistic surrounds (68). The explanation for the latter finding was that Cx36 gap junctions among a variety of amacrine cells, or among ganglion cells and amacrine cells, contribute to the formation of surrounds (68), again supporting inner retinal regulation of ganglion cell spatial response properties and confirming a role for Cx36 in spatial tuning. The reduction in photopic visual acuity in Cx36-DEDD homozygous mice indicates that plasticity of Cx36 electrical synapses is required to tune ganglion cell spatial response profiles optimally.

Although we have initially used the Cx36-DEDD mouse to investigate normal retinal functions, there are many potential uses for this model system in investigation of neurological disorders. There is evidence to suggest that unusually high electrical coupling occurs in certain pathological conditions. In mice with inherited retinal degeneration, retinal ganglion cells are hyperactive (69–71). This has been attributed to oscillations in the AII amacrine cell-bipolar cell network, in part due to excessive gap junction coupling that can be suppressed by knocking out Cx36 or administering dopamine receptor agonists to reduce coupling (72–74). Furthermore, AII amacrine cell gap junctions in rd models show excessive Cx36 phosphorylation (75), in keeping with these observations. More generally, excessive neuronal gap junction coupling has been found to occur in a variety of traumatic and ischemic brain injury models, contributing to neuronal cell death (76). The Cx36-DEDD mouse provides a new model system to study the contributions of neuronal coupling to neuron loss in neurodegenerative diseases and injuries.

## Materials and Methods

### Clones for in vitro expression studies

Clones for in vitro expression studies were developed using a mouse Cx36 cDNA clone with a C-terminal EGFP tag (42). Mutations were introduced using the QuickChange Multisite mutagenesis kit (Agilent, Santa Clara, CA) following the manufacturer’s instructions. Primers used for mutagenesis are listed in Table 1.

**Table 1.**
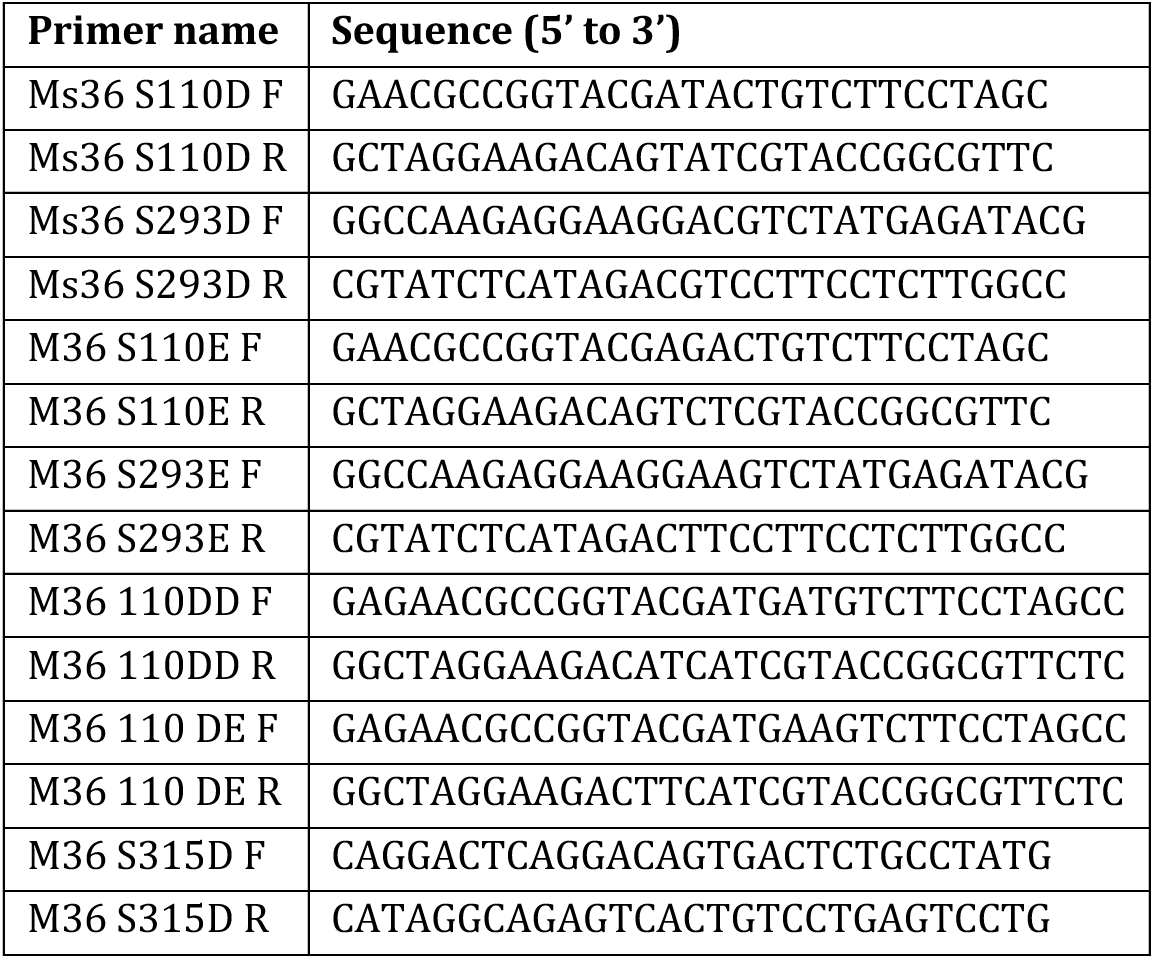
Primers used for development of phosphomimetic mutants.

Mutations were introduced into Cx36 in a stepwise manner and mutant constructs tested for functional regulation as described in the Results and Figure 1. All clones were verified by DNA sequencing. Cx36-EGFP and all mutant constructs will be made available through Addgene [authors will update this statement with clone numbers when Addgene deposit is complete].

### Tracer coupling studies in cultured cells

Regulation and magnitude of coupling supported by Cx36 mutants was measured by tracer coupling in transfected HeLa cells (catalog #CCL2, ATCC, Manassas, VA). Cells were grown in MEM with essential amino acids + sodium pyruvate, 10% FBS and Penicillin/Streptomycin/Fungizone. Cell culture reagents were obtained from Gibco/ThermoFisher (Gaithersburg, MD). Cells were plated onto tissue culture coated coverslips (ThermoFisher, Waltham, MA), grown to 70-80% confluency, and transiently transfected with 2 µg of DNA per dish using GenePorter 2 (Genlantis, San Diego, CA) following the manufacturer’s protocol.

Gap junction coupling was analyzed by measuring tracer diffusion of Neurobiotin (Vector Laboratories, Burlingame, CA) loaded into cells by scrape loading (31). Cells were incubated in cell maintaining solution (CMS - containing 150 mM NaCl, 6.2 mM KCl, 1. 2mM NaH_2_PO_4_, 1.2 mM MgSO_4_, 2. 5mM CaCl_2_, 10 mM glucose, 10 mM HEPES pH 7.4) either alone (control) or containing 10 µM protein kinase A antagonist Rp-8-cpt-cAMPs or 10 µM protein kinase A agonist Sp-8-cpt-cAMPs (both from Axxora LLC, Farmingdale, NY) for 10 min at 35°C. Fresh solutions of appropriate drug were added to each dish along with 0.1% Neurobiotin. Cells were scraped with a 26-gauge needle, allowed to incubate 10 min, rinsed three times in CMS, and fixed for 1 hr with 4% (w/v) formaldehyde (PFA; Electron Microscopy Sciences, Hatfield, PA) in 0.1 M phosphate buffer (PB; pH 7.4). Fixed cells were rinsed briefly in 0.1 M PB, permeabilized with phosphate buffered saline, pH 7.4 (PBS), 0.1% Triton X-100, 0.1% Na azide (PBSTA) for 1 hr, and labeled with Cy3-strepavidin (1:500; Jackson ImmunoResearch, West Valley, PA) for 1.5 hr. Coverslips were washed with PBSTA for 1 hr and mounted on slides using Vectashield mounting medium (Vector Laboratories) and imaged with a fluorescence microscope at 40x magnification.

In each experiment, a minimum of five loaded regions along the scraped edge of the cells was imaged for measurement. Images were collected using HCImage software (Hamamatsu Photonics, Sewickley, PA) or Micro-Manager (77) and analyzed with SimplePCI (Hamamatsu) and Matlab (MathWorks, Natick, MA) software. Intensity of Neurobiotin/Cy3-streptavidin signal was measured in 2 µm circles, with the brightest regions of individual cells selected. Cell-to-cell distance was measured from center to center of adjacent cells. Tracer diffusion was estimated by fitting data from groups of cells extending out from a loaded cell along the scraped edge of the coverslip using a linear compartmental diffusion model implemented by Mills and Massey (56, 78). This model assumes the cells are arranged in a linear chain connected by Cx36 gap junctions that are characterized by a rate-limiting diffusion coefficient k. Independent measurements of k were made for each of the 5 to 8 loaded regions examined within a single experiment. Three to six experiments were performed and all values of k were used for comparisons. Statistical comparisons were performed with Prism (GraphPad Software, San Diego, CA).

### Development of Cx36 targeting construct

Genomic elements of the mouse *Cx36* (*gjd2*) gene were obtained from genomic Bac clone RP23-230H3 (BacPac Resources, Emeryville, CA), derived from female C57BL/6J mice. The targeting construct was assembled with a 5’ targeting arm containing 3 kb of the *Cx36* gene including exon 1 and most of intron 1, and a 3’ targeting arm including 5.8 kb beginning at the end of the *Cx36* open reading frame (see Fig 2A). A LoxP site was inserted 88 bp before the wild type exon 2, and an artificial polyadenylation signal was inserted 1 kb after the *Cx36* exon 2 open reading frame.

This was followed by a self-deleting Neomycin selection cassette, which was deleted in the targeted embryonic stem (ES) cell lines and is not contained within the targeted animals. These elements were followed by the second LoxP site, the *Cx36-DEDD* mutant exon 2, an internal ribosome entry sequence (IRES) and the coding sequence of tdTomato (Fig 2A). Final construction of the targeting construct, as well as targeting and selection of ES cell lines, was performed by contract with Cyagen Biosciences (Santa Clara, CA).

A preliminary construct containing a short fragment of the *Cx36* promoter and exon 1, derived from a previous Cx36 transgene construct (79), a loxP site in intron 1, and the Cx36-DEDD mutant exon 2 with IRES dsRed2, was developed to test mRNA splicing across the LoxP site, IRES function and expression of the mutant Cx36. This construct was transferred into the EGFP-N1 vector backbone containing a CMV promoter to drive generalized expression in mammalian cells. Transfection of this construct into Hela cells produced cytoplasmic dsRed expression and Cx36 protein detectable by immunostaining at gap junctions (data not shown), confirming that splicing and IRES function were successful.

### Mice

Care and use of experimental animals was performed in accordance with the ARVO statement for the use of animals in ophthalmic and vision research and US Public Health Service guidelines, and were approved by the Institutional Animal Care and Use Committees at the University of Texas Health Science Center at Houston and the University of Houston. Connexin 36 phosphomimetic (Cx36-DEDD) conditional knockin mice in C57BL/6 background were custom generated through Cyagen Biosciences.

Two founder strains derived from two independent ES cell clones were obtained. One strain, termed “A+B” was used throughout this study. Wild type (WT) C57BL/6J mice were from Jackson Laboratory (Bar Harbor, ME; IMSR Cat# JAX:000664, RRID:IMSR_JAX:000664). Six3-Cre transgenic mice (48) were a gift from Yasuhide Furuta, MD Anderson Cancer Center. Strains were crossed as required to obtain genotypes used in this study. Animals were genotyped by PCR amplification of tail snip DNA. PCR primers used for genotyping are listed in Table 2. Both male and female mice were used without preference throughout this study.

**Table 2.**
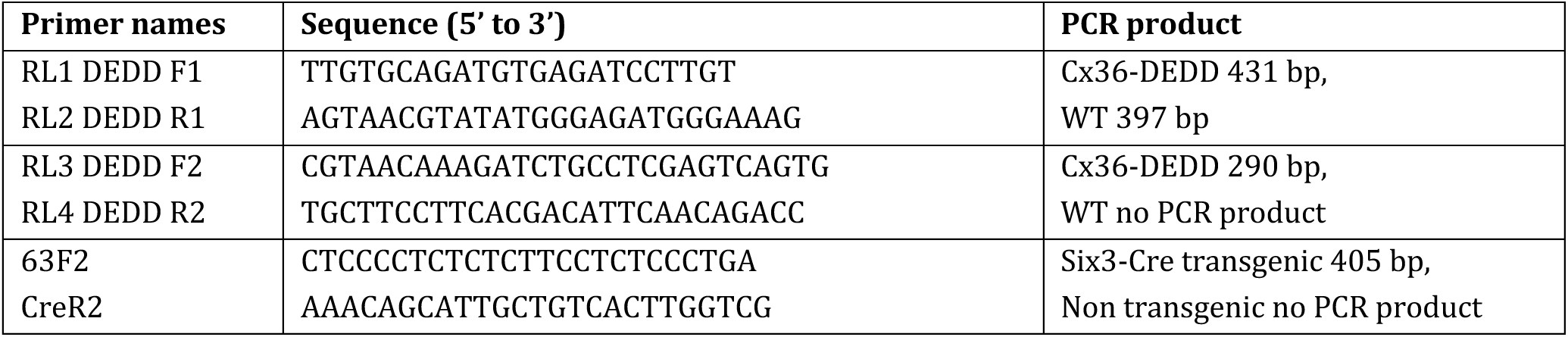
Primers used to genotype Cx36-DEDD mice.

### Immunostaining

Mice were anesthetized with isoflurane and sacrificed by cervical dislocation and eyes were quickly dissected in PBS and lenses removed. The eyecup was fixed at room temperature (RT) for 30 min with 2% PFA in PBS. Fixed tissue was cryoprotected in 30% sucrose in PBS overnight at 4°C and embedded in OCT (Sakura Finetek, Torrance, CA, USA). Tissue was sectioned at 12 µm thickness using a cryostat and blocked with 10% normal donkey serum (Jackson ImmunoResearch) in PBSTA at RT for 30 min.

Primary antibodies used were mouse anti-Cx36 (1:500; MAB3045, RRID:AB_94632, EMD Millipore, Billerica, MA), mouse anti-Cx36 (1:250; 37-4600, RRID:AB_2533320, Thermofisher, Rockford, IL, USA) and rabbit anti-RFP (1:200; ab62341, RRID:AB_945213, Abcam, Waltham, MA). Antibodies were diluted in PBSTA plus 5% donkey serum and applied to sections overnight at RT. Sections were washed extensively with PBSTA and incubated with Alexa Fluor 488 or Cy3-labeled secondary antibodies made in donkey (Jackson ImmunoResearch) at 1:500 dilution in PBSTA. The secondary antibodies were incubated for 1 hr at RT and then washed extensively with PBS. Nuclei were counterstained by mounting with Vectashield mounting medium with DAPI (Vector Laboratories). Samples were imaged on a Zeiss LSM 800 confocal microscope with 40x/1.4 NA objective. Imaging settings were conserved between samples from animals of different genotypes. Images were mapped into modified color space and adjusted for brightness using Fiji (ImageJ2 version 2.14.0). Where made, brightness adjustments were made to all samples equally.

### Retinal tracer coupling measurements

All experiments were performed on light-adapted animals in the daytime phase of their light cycle. Mice were anesthetized with isoflurane and sacrificed by cervical dislocation and eyes were quickly dissected in oxygenated Ames’ medium (Sigma-Aldrich, St. Louis, MO) supplemented with 26 mM NaHCO_3_, pH 7.4 at 35°C. The retinas were removed and flat mounted onto nitrocellulose filter paper (EMD Millipore) with photoreceptor side down. The flat mount retina was incubated in Ames’ medium equilibrated with 95% O_2_ and 5% CO_2_ for 20 min with DAPI to label cell nuclei. Sharp electrodes were pulled from thin-wall borosilicate glass (Sutter Instrument, Novato, CA) to 20 MΩ resistance and tip-filled with 4% Neurobiotin (Vector Laboratories) and 0.5% Lucifer Yellow CH (ThermoFisher) in 0.1 M PB, pH 7.4. Electrodes were back-filled with 3M LiCl. AII amacrine cells were injected by iontophoresis (±1 nA, 3 Hz) for 3-5 minutes and tracer was allowed to diffuse for at least 20 min, during which time the retinas were continuously superfused with O_2_/CO_2_-bubbled Ames’ medium at 35°C. Several injections were performed per retina, and precise injection times were recorded to enable modeling of tracer diffusion, which requires a measure of diffusion time (56).

After tracer injection and diffusion periods, the retinas were fixed for 30 minutes in 4% PFA in 0.1 M PB. Tracer-coupled networks were visualized by incubation with Alexa-488 Streptavidin (1:200; Jackson ImmunoResearch) in 0.1 M PB overnight at RT. Whole-mount retinas were then removed from the filter paper backing, mounted on microscope slides, and imaged with a confocal microscope. Coupling among AII amacrine cells was quantified as the diffusion coefficient of the Neurobiotin tracer, which was calculated by fitting the fluorescence intensities of AII somata and distance from the injected cell to a linear compartmental diffusion model as described above for tracer coupling in cell culture (56, 78). Statistical comparisons were performed with GraphPad Prism.

### Paired rod–cone patch-clamp recordings

Adult mice (2–6 months of age, either sex) were housed in a 12 h light/12 h dark cycle (lights on at 07:00) for at least 2 weeks before experiments. Mice were dark-adapted overnight, anesthetized with ketamine/xylazine (100/10 mg·kg⁻¹, intramuscular) and one eye was rapidly enucleated into oxygenated Ames’ medium supplemented with 23 mM NaHCO₃ (Sigma-Aldrich). Manipulations of the eyes and retina were carried out under infrared illumination using night-vision goggles to maintain deep dark adaptation unless otherwise stated (26, 27).

### Retinal slice preparation and solutions

The neural retina was isolated from the eyecup under a dissecting microscope by removing the cornea and lens and gently peeling away the sclera and retinal pigment epithelium, after which the retina was transferred ganglion-cell side down onto 0.45 µm HAWP filter paper (Millipore) submerged in Ames’ medium. Retinal slices (∼200 µm thick) were cut on a razor-blade tissue chopper together with the filter paper and rotated by 90° in a custom recording chamber so that all layers were exposed, then secured to the glass bottom with small bands of liquid silicone to minimize movement. Slices were superfused at 2 ml·min⁻¹ with bicarbonate-buffered Ames’ solution maintained at 32°C and bubbled continuously with 5% CO₂/95% O₂ and allowed to recover for ≥30 min before recordings; slices remained submerged and were never exposed to air. Patch pipettes contained an intracellular solution composed of (in mM): 10 KCl, 120 K-D-gluconate, 2 MgCl₂, 5 Na₂-ATP, and 1 Na₃-GTP, adjusted to pH 7.25 with KOH and ∼275 mosmol·l⁻¹; electrodes were backfilled with this solution containing 25 µM β-escin as perforating agent. DEDD littermate control and mutant slices were recorded under drug-free control conditions.

### Electrode fabrication and recording configuration

Patch pipettes were pulled from borosilicate glass capillaries (OD 1.2 mm, ID 0.69 mm; Sutter Instruments or WPI) on a P-1000 puller to yield a short shank and rapid taper; pipette resistance in bath solution was typically ∼15 MΩ. Two identical electrodes were mounted on independent MP-285 micromanipulators on a fixed-stage Olympus BX51WI microscope equipped with infrared (>900 nm) differential interference contrast optics and connected to separate patch-clamp amplifiers (Axopatch 200B and Dagan 3900A) whose outputs were digitized with a Digidata interface and recorded using pClamp; signals were filtered at 2 kHz and sampled at 10 kHz. Positive pressure was applied during approach to the slice to keep electrode tips clean, and gigaseals (>1 GΩ) were formed on targeted photoreceptors before β-escin–mediated perforation; series resistance after perforation was maintained between 20 and 33 MΩ, and cells with larger access resistance or unstable holding currents were excluded.

### Identification of rods and cones and pair selection

Under infrared DIC, rods were targeted in the middle of the outer nuclear layer (ONL), where somata are exclusively rod, whereas cone pedicles were identified as highly contrasted, enlarged terminals at the base of the ONL and within the outer plexiform layer. For rod–cone paired recordings, one electrode was positioned on a rod soma within the rod-only region and the second electrode was positioned on the pedicle of an adjacent cone at the corresponding position in the OPL, such that the two cells were nearest neighbors in the mosaic. In a subset of experiments, Lucifer Yellow was included in the pipette solution to confirm cell identity and relative position post hoc by confocal imaging of the filled rod and cone and the surrounding photoreceptor array.

### Voltage-clamp protocols and junctional conductance measurements

After stable perforated patch configuration was obtained in both cells, the cone designated as the “slave” was held at −35 mV, while the “driver” cell was stepped with 50-ms voltage pulses from −50 to +50 mV in 10-mV increments to evoke transjunctional currents. Junctional currents were measured in the slave cell as the steady current obtained by averaging data between 20 and 30 ms after the onset of each voltage step, a window chosen to minimize contamination by voltage-gated conductances such as hyperpolarization-activated cation current and calcium-activated chloride current. For each pair, the junctional current was plotted as a function of the transjunctional voltage, and the junctional conductance (Gⱼ) was computed as the slope of the best-fit linear regression; pairs displaying nonlinear I–V relationships or obvious contamination by voltage-dependent currents were discarded. During paired recordings, no light stimuli were delivered to avoid acute modulation of gap-junction conductance or bleaching of pigment, and the chamber was enclosed in a light-tight Faraday cage lined with conductive plastic to block ambient light and reduce electrical noise. The liquid junction potential (∼10 mV) was not corrected, and all voltages are reported as command potentials relative to the bath.

### Noise estimation, background subtraction, and inclusion criteria

Because junctional conductances between mammalian photoreceptors are small (tens to hundreds of pS), background noise was measured repeatedly with both electrodes in the bath at similar separation distances but not sealed to cells, and the resulting ohmic current–voltage relation (typically ∼100 pS) was subtracted from all cell-pair measurements. Recordings exhibiting high-frequency baseline fluctuations or obvious amplifier drift were terminated, and Ag/AgCl bridges were re-chlorinated when necessary to reduce noise. Only pairs with stable access resistance, linear I–V relations, and junctional conductance values above the noise-limited detection threshold (∼100 pS) were included in the analysis of rod–cone coupling in DEDD littermate control and mutant mice.

### Optomotor response measurements

Image forming visual function was examined by observing the optomotor reflex with an Optodrum system (Striatech, Tubingen, Germany; Software version 1.5.3). Briefly, a mouse was placed on a platform without movement restriction surrounded by a virtual cylinder of vertical gratings displayed by four computer monitors. The position of the animal’s head was continuously tracked in real time by Optodrum software. Analysis of the head movement, in combination with the parameters of the grating, were evaluated to decide if an optomotor reflex was induced.

The grating stimuli were presented at a fixed drifting speed of 12 degrees/sec and fixed mean luminance of 70 cd/m^2^ (illuminance at the platform = 82 lux). Visual acuity was assessed by varying spatial frequency (i.e. the number of black/white cycles within 1 visual degree) at fixed 99.7% contrast. Contrast sensitivity was assessed by varying stripe contrast (i.e. brightness difference between the white and black stripes) at a spatial frequency of 0.1 cycle/degree. The software uses a staircase algorithm to determine the parameters of the next stimulus, and repeats tests until a threshold is determined. Both types of measurements were performed with both clockwise rotation of the stimuli (testing predominantly left eye function) and counterclockwise rotation (testing predominantly right eye function). These were done sequentially in the same test session in our experiments. Almost all animals were tested on two different occasions, with test results from the two directions of movement and the two testing sessions averaged. The animals were allowed at least one day between tests. All experiments were performed on light-adapted animals in the daytime phase of their light cycle, between 10:00 am and 12:00 pm. 4- to 6-month-old mice, either phosphomimetic Connexin 36 knockin or their littermate controls were used. Statistical comparisons were performed with GraphPad Prism.

## Acknowledgments

The authors would like to thank Dr. Rane Liu from Cyagen Biosciences for expert services and coordination of the conditional knock-in mouse project. We would like to thank Dr. Eva Zsigmond and Aleksey Domozhirov from the University of Texas Health Science Center-Houston, Transgenic and Stem Cells Service Unit for expert services performing the original knock-in projects.

This research was supported by NIH grants R01EY012857 and R21EY027965 (JO), R01EY024376 (C-AM), R01EY032508 (CPR), core grants P30EY028102 and P30EY007551, and the Louisa Stude Sarofim endowment (JO).

## Author Contributions

**Y-P.L.** performed research, contributed new reagents or analytic tools, analyzed data; **C.K.M.** performed research, analyzed data; **N.J.** performed research, analyzed data; **Z.Z.** performed research, analyzed data; **A.N.** performed research, analyzed data; **N.A.B.** performed research, analyzed data; **S.T.** performed research, analyzed data; **Y.L.** performed research, analyzed data; **H.L.** performed research, contributed new reagents or analytic tools, analyzed data; **S.L.** performed research, analyzed data; **C-A.M.** designed research, acquired funding; **S.W.W.** designed research; **C.P.R.** designed research, analyzed data, acquired funding; **J.O.** designed research, contributed new reagents or analytic tools, analyzed data, acquired funding, wrote the paper. All authors edited and revised the manuscript.

