## Supplemental for "Retinal electrical synapse plasticity is required for optimal visual performance"

**This PDF file includes:**

Figures S1 to S2
Tables S1 to S2

**Figures**

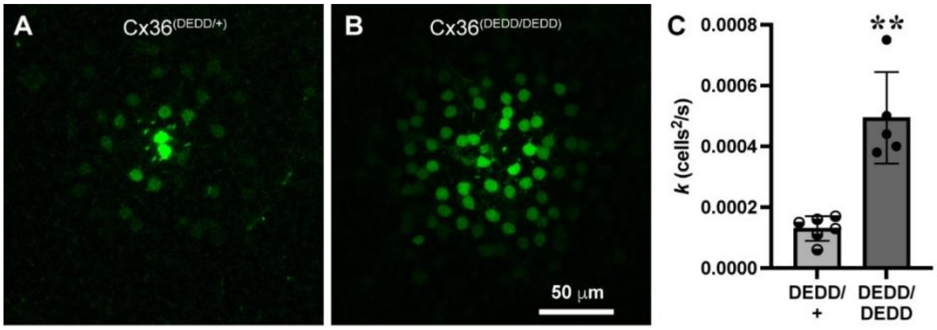

**Supplemental Figure 1. Tracer coupling in AII amacrine cells of Cx36-DEDD mice.** **A**. Neurobiotin tracer injection into a single AII amacrine cell in Six3-Cre/Cx36$^{(DEDD/+)}$ heterozygous mouse retina. **B**. Neurobiotin tracer injection into a single AII amacrine cell in Six3-Cre/Cx36$^{(DEDD/DEDD)}$ homozygous mouse retina. Scale bar in B applies to A as well. **C**. Diffusion coefficients for Neurobiotin tracer coupling in Six3-Cre/Cx36$^{(DEDD/+)}$ and Six3-Cre/Cx36$^{(DEDD/DEDD)}$ mouse retina. Bars are means +/- SD. ** $p < 0.01$.

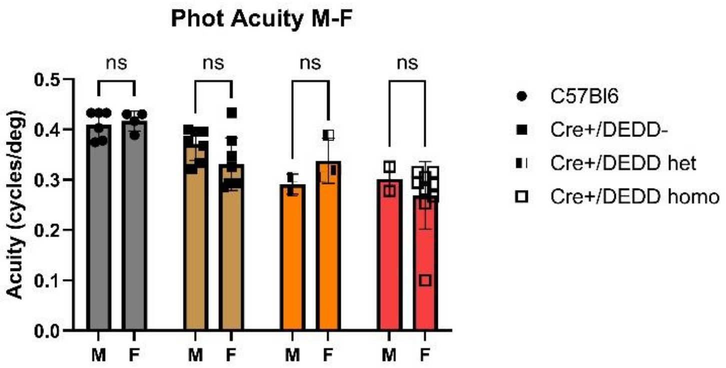

**Supplemental Figure 2**. **Effect of sex on photopic visual acuity at 99.7% contrast.** There were no significant differences between female and male mice of any genotype tested in this parameter.

**Tables**

**Table S1.** Statistical comparisons of tracer coupling in vitro.

| **Figure 1C** | | | | | | | |
|---|---|---|---|---|---|---|---|
| Ordinary two-way ANOVA | | | | | | | |
| Alpha = 0.05 | | | | | | | |
| Source of Variation | % of total variation | P value | P value summary | Significant? | | | |
| Interaction | 20.22 | <0.0001 | **** | Yes | | | |
| PKA | 24.92 | <0.0001 | **** | Yes | | | |
| Mutation | 16.65 | <0.0001 | **** | Yes | | | |
| ANOVA table | SS (Type III) | DF | MS | F (DFn, DFd) | P value | | |
| Interaction | 1.22E-05 | 6 | 2.04E-06 | F (6, 164) = 14.59 | P<0.0001 | | |
| PKA | 1.51E-05 | 2 | 7.53E-06 | F (2, 164) = 53.95 | P<0.0001 | | |
| Mutation | 1.01E-05 | 3 | 3.36E-06 | F (3, 164) = 24.03 | P<0.0001 | | |
| Residual | 2.29E-05 | 164 | 1.40E-07 | | | | |
| Total | 6.05E-05 | 175 | | | | | |
| Tukey's multiple comparisons test | Predicted (LS) Mean diff. | 95.00% CI of diff. | Below threshold? | Summary | Adjusted P Value | N1 | N2 |
| con | | | | | | | |
| EV vs. WT | -0.0001553 | -0.0004704 to 0.0001598 | No | ns | 0.5772 | 20 | 18 |
| EV vs. DTD | -0.0001445 | -0.0004986 to 0.0002096 | No | ns | 0.7148 | 20 | 12 |
| EV vs. ETE | -0.0001388 | -0.0005029 to 0.0002252 | No | ns | 0.7555 | 20 | 11 |
| WT vs. DTD | 1.08E-05 | -0.0003506 to 0.0003723 | No | ns | 0.9998 | 18 | 12 |
| WT vs. ETE | 1.65E-05 | -0.0003546 to 0.0003877 | No | ns | 0.9994 | 18 | 11 |
| DTD vs. ETE | 5.68E-06 | -0.0003991 to 0.0004105 | No | ns | >0.9999 | 12 | 11 |

| | | | | | | | |
|---|---|---|---|---|---|---|---|
| Rp | | | | | | | |
| EV vs. WT | -0.001404 | -0.001732 to -0.001076 | Yes | **** | <0.0001 | 18 | 17 |
| EV vs. DTD | -0.001269 | -0.001622 to -0.0009164 | Yes | **** | <0.0001 | 18 | 13 |
| EV vs. ETE | -0.0004656 | -0.0008367 to -9.439e-005 | Yes | ** | 0.0074 | 18 | 11 |
| WT vs. DTD | 0.0001344 | -0.0002229 to 0.0004917 | No | ns | 0.7632 | 17 | 13 |
| WT vs. ETE | 0.0009382 | 0.0005630 to 0.001314 | Yes | **** | <0.0001 | 17 | 11 |
| DTD vs. ETE | 0.0008038 | 0.0004065 to 0.001201 | Yes | **** | <0.0001 | 13 | 11 |
| Sp | | | | | | | |
| EV vs. WT | -0.0001726 | -0.0005058 to 0.0001607 | No | ns | 0.5362 | 16 | 18 |
| EV vs. DTD | -7.67E-06 | -0.0003875 to 0.0003722 | No | ns | >0.9999 | 16 | 11 |
| EV vs. ETE | -0.0001031 | -0.0004830 to 0.0002767 | No | ns | 0.8951 | 16 | 11 |
| WT vs. DTD | 0.0001649 | -0.0002063 to 0.0005361 | No | ns | 0.6572 | 18 | 11 |
| WT vs. ETE | 6.94E-05 | -0.0003017 to 0.0004406 | No | ns | 0.9622 | 18 | 11 |
| DTD vs. ETE | -9.55E-05 | -0.0005090 to 0.0003181 | No | ns | 0.9322 | 11 | 11 |
| EV | | | | | | | |
| con vs. Rp | 5.86E-05 | -0.0002286 to 0.0003457 | No | ns | 0.8798 | 20 | 18 |
| con vs. Sp | -3.88E-06 | -0.0003003 to 0.0002925 | No | ns | 0.9995 | 20 | 16 |
| Rp vs. Sp | -6.24E-05 | -0.0003661 to 0.0002412 | No | ns | 0.8779 | 18 | 16 |
| WT | | | | | | | |
| con vs. Rp | -0.00119 | -0.001489 to -0.0008910 | Yes | **** | <0.0001 | 18 | 17 |
| con vs. Sp | -2.11E-05 | -0.0003157 to 0.0002735 | No | ns | 0.9843 | 18 | 18 |
| Rp vs. Sp | 0.001169 | 0.0008699 to 0.001468 | Yes | **** | <0.0001 | 17 | 18 |
| DTD | | | | | | | |

| | | | | | | | |
|---|---|---|---|---|---|---|---|
| con vs. Rp | -0.001066 | -0.001420 to -0.0007126 | Yes | **** | <0.0001 | 12 | 13 |
| con vs. Sp | 0.000133 | -0.0002359 to 0.0005019 | No | ns | 0.671 | 12 | 11 |
| Rp vs. Sp | 0.001199 | 0.0008373 to 0.001561 | Yes | **** | <0.0001 | 13 | 11 |
| ETE | | | | | | | |
| con vs. Rp | -0.0002682 | -0.0006450 to 0.0001087 | No | ns | 0.2147 | 11 | 11 |
| con vs. Sp | 3.18E-05 | -0.0003450 to 0.0004087 | No | ns | 0.9783 | 11 | 11 |
| Rp vs. Sp | 0.0003 | -7.683e-005 to 0.0006768 | No | ns | 0.1469 | 11 | 11 |
| **Figure 1D** | | | | | | | |
| Ordinary two-way ANOVA | | | | | | | |
| Alpha = 0.05 | | | | | | | |
| Source of Variation | % of total variation | P value | P value summary | Significant? | | | |
| Interaction | 18.55 | <0.0001 | **** | Yes | | | |
| PKA | 36.08 | <0.0001 | **** | Yes | | | |
| Mutation | 14.93 | <0.0001 | **** | Yes | | | |
| ANOVA table | SS (Type III) | DF | MS | F (DFn, DFd) | P value | | |
| Interaction | 1.51E-05 | 6 | 2.52E-06 | F (6, 167) = 13.81 | P<0.0001 | | |
| PKA | 2.94E-05 | 2 | 1.47E-05 | F (2, 167) = 80.57 | P<0.0001 | | |
| Mutation | 1.22E-05 | 3 | 4.05E-06 | F (3, 167) = 22.23 | P<0.0001 | | |
| Residual | 3.04E-05 | 167 | 1.82E-07 | | | | |
| Total | 8.14E-05 | 178 | | | | | |
| Tukey's multiple comparisons test | Predicted (LS) Mean diff. | 95.00% CI of diff. | Below threshold? | Summary | Adjusted P Value | N1 | N2 |
| con | | | | | | | |

| | | | | | | | |
|---|---|---|---|---|---|---|---|
| EV vs. WT | -0.0001553 | -0.0005153 to 0.0002046 | No | ns | 0.6778 | 20 | 18 |
| EV vs. DED | -3.78E-05 | -0.0004325 to 0.0003569 | No | ns | 0.9946 | 20 | 13 |
| EV vs. 3D | -0.0002147 | -0.0006094 to 0.0001800 | No | ns | 0.4939 | 20 | 13 |
| WT vs. DED | 0.0001176 | -0.0002857 to 0.0005208 | No | ns | 0.8737 | 18 | 13 |
| WT vs. 3D | -5.94E-05 | -0.0004626 to 0.0003439 | No | ns | 0.981 | 18 | 13 |
| DED vs. 3D | -0.0001769 | -0.0006115 to 0.0002576 | No | ns | 0.7163 | 13 | 13 |
| Rp | | | | | | | |
| EV vs. WT | -0.001404 | -0.001778 to -0.001029 | Yes | **** | <0.0001 | 18 | 17 |
| EV vs. DED | -0.0009489 | -0.001362 to -0.0005360 | Yes | **** | <0.0001 | 18 | 12 |
| EV vs. 3D | -0.001731 | -0.002168 to -0.001294 | Yes | **** | <0.0001 | 18 | 10 |
| WT vs. DED | 0.0004549 | 3.718e-005 to 0.0008726 | Yes | * | 0.0269 | 17 | 12 |
| WT vs. 3D | -0.0003268 | -0.0007683 to 0.0001148 | No | ns | 0.2234 | 17 | 10 |
| DED vs. 3D | -0.0007817 | -0.001256 to -0.0003073 | Yes | *** | 0.0002 | 12 | 10 |
| Sp | | | | | | | |
| EV vs. WT | -0.0001726 | -0.0005532 to 0.0002081 | No | ns | 0.6425 | 16 | 18 |
| EV vs. DED | 1.04E-06 | -0.0004220 to 0.0004241 | No | ns | >0.9999 | 16 | 12 |
| EV vs. 3D | 3.02E-05 | -0.0003929 to 0.0004533 | No | ns | 0.9977 | 16 | 12 |
| WT vs. DED | 0.0001736 | -0.0002393 to 0.0005865 | No | ns | 0.6954 | 18 | 12 |
| WT vs. 3D | 0.0002028 | -0.0002101 to 0.0006157 | No | ns | 0.5804 | 18 | 12 |
| DED vs. 3D | 2.92E-05 | -0.0004231 to 0.0004815 | No | ns | 0.9983 | 12 | 12 |
| EV | | | | | | | |
| con vs. Rp | 5.86E-05 | -0.0002695 to 0.0003866 | No | ns | 0.9065 | 20 | 18 |
| con vs. Sp | -3.88E-06 | -0.0003425 to 0.0003348 | No | ns | 0.9996 | 20 | 16 |
| Rp vs. Sp | -6.24E-05 | -0.0004093 to 0.0002845 | No | ns | 0.905 | 18 | 16 |

| | | | | | | | |
|---|---|---|---|---|---|---|---|
| WT | | | | | | | |
| con vs. Rp | -0.00119 | -0.001531 to -0.0008485 | Yes | **** | <0.0001 | 18 | 17 |
| con vs. Sp | -2.11E-05 | -0.0003576 to 0.0003154 | No | ns | 0.9879 | 18 | 18 |
| Rp vs. Sp | 0.001169 | 0.0008273 to 0.001510 | Yes | **** | <0.0001 | 17 | 18 |
| DED | | | | | | | |
| con vs. Rp | -0.0008526 | -0.001257 to -0.0004484 | Yes | **** | <0.0001 | 13 | 12 |
| con vs. Sp | 3.49E-05 | -0.0003692 to 0.0004391 | No | ns | 0.9772 | 13 | 12 |
| Rp vs. Sp | 0.0008875 | 0.0004753 to 0.001300 | Yes | **** | <0.0001 | 12 | 12 |
| 3D | | | | | | | |
| con vs. Rp | -0.001457 | -0.001882 to -0.001033 | Yes | **** | <0.0001 | 13 | 10 |
| con vs. Sp | 0.000241 | -0.0001631 to 0.0006452 | No | ns | 0.3379 | 13 | 12 |
| Rp vs. Sp | 0.001698 | 0.001266 to 0.002131 | Yes | **** | <0.0001 | 10 | 12 |
| **Figure 1E** | | | | | | | |
| Ordinary two-way ANOVA | | | | | | | |
| Alpha = 0.05 | | | | | | | |
| Source of Variation | % of total variation | P value | P value summary | Significant? | | | |
| Interaction | 19.69 | <0.0001 | **** | Yes | | | |
| PKA | 5.657 | 0.0007 | *** | Yes | | | |
| Mutation | 31.91 | <0.0001 | **** | Yes | | | |
| ANOVA table | SS (Type III) | DF | MS | F (DFn, DFd) | P value | | |
| Interaction | 1.64E-05 | 6 | 2.74E-06 | F (6, 117) = 8.902 | P<0.0001 | | |
| PKA | 4.72E-06 | 2 | 2.36E-06 | F (2, 117) = 7.673 | P=0.0007 | | |
| Mutation | 2.66E-05 | 3 | 8.87E-06 | F (3, 117) = 28.86 | P<0.0001 | | |
| Residual | 3.60E-05 | 117 | 3.08E-07 | | | | |

| | | | | | | | |
|---|---|---|---|---|---|---|---|
| Total | 8.34E-05 | 128 | | | | | |
| Tukey's multiple comparisons test | Predicted (LS) Mean diff. | 95.00% CI of diff. | Below threshold? | Summary | Adjusted P Value | N1 | N2 |
| con | | | | | | | |
| EV vs. WT | -4.55E-06 | -0.0006208 to 0.0006117 | No | ns | >0.9999 | 11 | 11 |
| EV vs. 315D | -0.0001864 | -0.0008026 to 0.0004299 | No | ns | 0.8597 | 11 | 11 |
| EV vs. DEDD | -0.001156 | -0.001759 to -0.0005528 | Yes | **** | <0.0001 | 11 | 12 |
| WT vs. 315D | -0.0001818 | -0.0007981 to 0.0004344 | No | ns | 0.8683 | 11 | 11 |
| WT vs. DEDD | -0.001152 | -0.001755 to -0.0005482 | Yes | **** | <0.0001 | 11 | 12 |
| 315D vs. DEDD | -0.0009697 | -0.001573 to -0.0003664 | Yes | *** | 0.0003 | 11 | 12 |
| Rp | | | | | | | |
| EV vs. WT | -0.00119 | -0.001836 to -0.0005437 | Yes | **** | <0.0001 | 10 | 10 |
| EV vs. 315D | -0.0006458 | -0.001265 to -2.703e-005 | Yes | * | 0.0372 | 10 | 12 |
| EV vs. DEDD | -0.0009932 | -0.001625 to -0.0003617 | Yes | *** | 0.0004 | 10 | 11 |
| WT vs. 315D | 0.0005442 | -7.464e-005 to 0.001163 | No | ns | 0.1058 | 10 | 12 |
| WT vs. DEDD | 0.0001968 | -0.0004346 to 0.0008283 | No | ns | 0.8485 | 10 | 11 |
| 315D vs. DEDD | -0.0003473 | -0.0009506 to 0.0002559 | No | ns | 0.4404 | 12 | 11 |
| Sp | | | | | | | |
| EV vs. WT | 2.80E-05 | -0.0006183 to 0.0006743 | No | ns | 0.9995 | 10 | 10 |
| EV vs. 315D | -0.001473 | -0.002105 to -0.0008417 | Yes | **** | <0.0001 | 10 | 11 |
| EV vs. DEDD | -0.00155 | -0.002196 to -0.0009037 | Yes | **** | <0.0001 | 10 | 10 |
| WT vs. 315D | -0.001501 | -0.002133 to -0.0008697 | Yes | **** | <0.0001 | 10 | 11 |
| WT vs. DEDD | -0.001578 | -0.002224 to -0.0009317 | Yes | **** | <0.0001 | 10 | 10 |
| 315D vs. DEDD | -7.68E-05 | -0.0007083 to 0.0005546 | No | ns | 0.9889 | 11 | 10 |

| | | | | | | | |
|---|---|---|---|---|---|---|---|
| EV | | | | | | | |
| con vs. Rp | 2.27E-06 | -0.0005729 to 0.0005774 | No | ns | >0.9999 | 11 | 10 |
| con vs. Sp | -1.77E-05 | -0.0005929 to 0.0005574 | No | ns | 0.9971 | 11 | 10 |
| Rp vs. Sp | -2.00E-05 | -0.0006087 to 0.0005687 | No | ns | 0.9964 | 10 | 10 |
| WT | | | | | | | |
| con vs. Rp | -0.001183 | -0.001758 to -0.0006080 | Yes | **** | <0.0001 | 11 | 10 |
| con vs. Sp | 1.48E-05 | -0.0005603 to 0.0005900 | No | ns | 0.9979 | 11 | 10 |
| Rp vs. Sp | 0.001198 | 0.0006093 to 0.001787 | Yes | **** | <0.0001 | 10 | 10 |
| 315D | | | | | | | |
| con vs. Rp | -0.0004572 | -0.001007 to 9.227e-005 | No | ns | 0.1229 | 11 | 12 |
| con vs. Sp | -0.001305 | -0.001866 to -0.0007433 | Yes | **** | <0.0001 | 11 | 11 |
| Rp vs. Sp | -0.0008473 | -0.001397 to -0.0002979 | Yes | ** | 0.0011 | 12 | 11 |
| DEDD | | | | | | | |
| con vs. Rp | 0.0001652 | -0.0003843 to 0.0007146 | No | ns | 0.756 | 12 | 11 |
| con vs. Sp | -0.0004117 | -0.0009753 to 0.0001520 | No | ns | 0.197 | 12 | 10 |
| Rp vs. Sp | -0.0005768 | -0.001152 to -1.671e-006 | Yes | * | 0.0492 | 11 | 10 |
| **Figure 1F** | | | | | | | |
| Ordinary two-way ANOVA | | | | | | | |
| Alpha = 0.05 | | | | | | | |
| Source of Variation | % of total variation | P value | P value summary | Significant? | | | |
| Interaction | 23.68 | <0.0001 | **** | Yes | | | |
| PKA | 34.42 | <0.0001 | **** | Yes | | | |
| DEDD | 6.721 | 0.0018 | ** | Yes | | | |

| ANOVA table | SS | DF | MS | F (DFn, DFd) | P value | | |
|---|---|---|---|---|---|---|---|
| Interaction | 6.87E-07 | 1 | 6.87E-07 | F (1, 56) = 37.70 | P<0.0001 | | |
| PKA | 9.99E-07 | 1 | 9.99E-07 | F (1, 56) = 54.80 | P<0.0001 | | |
| DEDD | 1.95E-07 | 1 | 1.95E-07 | F (1, 56) = 10.70 | P=0.0018 | | |
| Residual | 1.02E-06 | 56 | 1.82E-08 | | | | |
| Total | 2.90E-06 | 59 | | | | | |
| Uncorrected Fisher's LSD | Mean diff. | 95.00% CI of diff. | Below threshold? | Summary | Individual P Value | N1 | N2 |
| con | | | | | | | |
| Cx36 WT vs. Cx36-DEDD | -0.000328 | -0.0004267 to -0.0002293 | Yes | **** | <0.0001 | 15 | 15 |
| glut | | | | | | | |
| Cx36 WT vs. Cx36-DEDD | 0.0001 | 1.267e-006 to 0.0001987 | Yes | * | 0.0472 | 15 | 15 |
| Cx36 WT | | | | | | | |
| con vs. glut | -0.000472 | -0.0005707 to -0.0003733 | Yes | **** | <0.0001 | 15 | 15 |
| Cx36-DEDD | | | | | | | |
| con vs. glut | -4.40E-05 | -0.0001427 to 5.473e-005 | No | ns | 0.3758 | 15 | 15 |

**Table S2.** Statistical comparisons of Optomotor response data.

| **Supplemental Figure 2** | | | | | | | |
|---|---|---|---|---|---|---|---|
| Ordinary two-way ANOVA | | | | | | | |
| Alpha = 0.05 | | | | | | | |
| Source of Variation | % of total variation | P value | P value summary | Significant? | | | |
| Interaction | 4.579 | 0.2936 | ns | No | | | |
| Sex | 0.08078 | 0.7954 | ns | No | | | |
| Genotype | 38.77 | <0.0001 | **** | Yes | | | |
| ANOVA table | SS (Type III) | DF | MS | F (DFn, DFd) | P value | | |
| Interaction | 0.008398 | 3 | 0.002799 | F (3, 33) = 1.292 | P=0.2936 | | |
| Sex | 0.0001481 | 1 | 0.0001481 | F (1, 33) = 0.06836 | P=0.7954 | | |
| Genotype | 0.0711 | 3 | 0.0237 | F (3, 33) = 10.94 | P<0.0001 | | |
| Residual | 0.07152 | 33 | 0.002167 | | | | |
| Tukey's multiple comparisons test | Predicted (LS) Mean diff. | 95.00% CI of diff. | Below threshold? | Summary | Adjusted P Value | N1 | N2 |
| C57Bl6 | | | | | | | |
| M vs. F | -0.007583 | -0.06872 to 0.05355 | No | ns | 0.8023 | 6 | 4 |
| Cre+/DEDD- | | | | | | | |
| M vs. F | 0.03899 | -0.01003 to 0.08801 | No | ns | 0.1151 | 7 | 8 |
| Cre+/DEDD het | | | | | | | |

| | | | | | | | |
|---|---|---|---|---|---|---|---|
| M vs. F | -0.04637 | -0.1328 to 0.04009 | No | ns | 0.2831 | 2 | 3 |
| Cre+/DEDD homo | | | | | | | |
| M vs. F | 0.03273 | -0.04131 to 0.1068 | No | ns | 0.375 | 2 | 9 |
| **Figure 4E** | | | | | | | |
| One-way ANOVA | | | | | | | |
| Alpha = 0.05 | | | | | | | |
| ANOVA summary | | | | | | | |
| F | 15.36 | | | | | | |
| P value | <0.0001 | | | | | | |
| P value summary | **** | | | | | | |
| Significant diff. among means (P < 0.05)? | Yes | | | | | | |
| R squared | 0.5547 | | | | | | |
| Brown-Forsythe test | | | | | | | |
| F (DFn, DFd) | 0.7960 (3, 37) | | | | | | |
| P value | 0.504 | | | | | | |
| P value summary | ns | | | | | | |
| Are SDs significantly different (P < 0.05)? | No | | | | | | |
| Bartlett's test | | | | | | | |
| Bartlett's statistic (corrected) | 7.091 | | | | | | |
| P value | 0.069 | | | | | | |
| P value summary | ns | | | | | | |

| | | | | | | | |
|---|---|---|---|---|---|---|---|
| Are SDs significantly different (P < 0.05)? | No | | | | | | |
| ANOVA table | SS | DF | MS | F (DFn, DFd) | P value | | |
| Treatment (between columns) | 0.1017 | 3 | 0.03391 | F (3, 37) = 15.36 | P<0.0001 | | |
| Residual (within columns) | 0.08167 | 37 | 0.002207 | | | | |
| Total | 0.1834 | 40 | | | | | |
| Tukey's multiple comparisons test | Mean diff. | 95.00% CI of diff. | Below threshold? | Summary | Adjusted P Value | N1 | N2 |
| C57Bl6 vs. Cre+/DEDD- | 0.06257 | 0.01098 to 0.1142 | Yes | * | 0.0122 | 10 | 15 |
| C57Bl6 vs. Cre+/DEDD het | 0.09275 | 0.02354 to 0.1620 | Yes | ** | 0.0049 | 10 | 5 |
| C57Bl6 vs. Cre+/DEDD homo | 0.137 | 0.08176 to 0.1922 | Yes | **** | <0.0001 | 10 | 11 |
| Cre+/DEDD- vs. Cre+/DEDD het | 0.03018 | -0.03507 to 0.09544 | No | ns | 0.6034 | 15 | 5 |
| Cre+/DEDD- vs. Cre+/DEDD homo | 0.07441 | 0.02425 to 0.1246 | Yes | ** | 0.0016 | 15 | 11 |
| Cre+/DEDD het vs. Cre+/DEDD homo | 0.04423 | -0.02393 to 0.1124 | No | ns | 0.3155 | 5 | 11 |
| **Figure 4F** | | | | | | | |
| One-way ANOVA | | | | | | | |
| Alpha = 0.05 | | | | | | | |
| ANOVA summary | | | | | | | |
| F | 24.79 | | | | | | |
| P value | <0.0001 | | | | | | |
| P value summary | **** | | | | | | |
| Significant diff. among means (P < 0.05)? | Yes | | | | | | |

| | | | | | | | |
|---|---|---|---|---|---|---|---|
| R squared | 0.6502 | | | | | | |
| Brown-Forsythe test | | | | | | | |
| F (DFn, DFd) | 0.7108 (3, 40) | | | | | | |
| P value | 0.5513 | | | | | | |
| P value summary | ns | | | | | | |
| Are SDs significantly different (P < 0.05)? | No | | | | | | |
| Bartlett's test | | | | | | | |
| Bartlett's statistic (corrected) | 3.288 | | | | | | |
| P value | 0.3494 | | | | | | |
| P value summary | ns | | | | | | |
| Are SDs significantly different (P < 0.05)? | No | | | | | | |
| ANOVA table | SS | DF | MS | F (DFn, DFd) | P value | | |
| Treatment (between columns) | 1179 | 3 | 393 | F (3, 40) = 24.79 | P<0.0001 | | |
| Residual (within columns) | 634.1 | 40 | 15.85 | | | | |
| Total | 1813 | 43 | | | | | |
| Tukey's multiple comparisons test | Mean diff. | 95.00% CI of diff. | Below threshold? | Summary | Adjusted P Value | N1 | N2 |
| C57Bl6 vs. Cre+/DEDD- | 3.435 | -0.9221 to 7.792 | No | ns | 0.1664 | 10 | 15 |
| C57Bl6 vs. Cre+/DEDD het | 1.812 | -4.033 to 7.658 | No | ns | 0.8394 | 10 | 5 |
| C57Bl6 vs. Cre+/DEDD homo | 12.79 | 8.374 to 17.21 | Yes | **** | <0.0001 | 10 | 14 |

| | | | | | | | |
|---|---|---|---|---|---|---|---|
| Cre+/DEDD- vs. Cre+/DEDD het | -1.622 | -7.134 to 3.889 | No | ns | 0.859 | 15 | 5 |
| Cre+/DEDD- vs. Cre+/DEDD homo | 9.358 | 5.392 to 13.32 | Yes | **** | <0.0001 | 15 | 14 |
| Cre+/DEDD het vs. Cre+/DEDD homo | 10.98 | 5.420 to 16.54 | Yes | **** | <0.0001 | 5 | 14 |
